# A conserved hydrophobic interaction governs GPCR–transducer association

**DOI:** 10.64898/2026.02.25.707970

**Authors:** Hyunggu Hahn, Emmanuel Flores-Espinoza, Anthony Nguyen, Martin Jung, Bianca Plouffe, Alex R. B. Thomsen

## Abstract

A central feature of G protein-coupled receptor (GPCR) desensitization is the direct competition between heterotrimeric G proteins and β-arrestins (βarrs) for an overlapping binding site within the intracellular receptor cavity. Although numerous high-resolution structures of GPCR–transducer complexes exist, the exact nature of this shared site and the molecular basis of transducer competition remain unclear. To investigate this, we employed an interdisciplinary approach integrating systemic mutational mapping, bioinformatics, and structural analysis across multiple classes of GPCRs and their transducers and regulators. We identified two highly conserved leucine residues within both the βarr finger loop and the Gα C-terminal α-helix, which engage a hydrophobic patch on GPCRs formed by TM3, TM5, and TM6 in a nearly identical manner, thereby stabilizing the complexes. Notably, the GPCR kinase N-terminal α-helix also contains hydrophobic residues that associate with this same receptor patch and are vital for the GPCR-GRK engagement. Our findings reveal a conserved hydrophobic interface that mediates direct competition among GPCR transducers and regulators suggesting a universal mechanism that governs receptor access and desensitization.

## Introduction

G protein-coupled receptors (GPCRs) are the largest family of transmembrane receptors with over 800 members that are involved in a wide range of physiological processes. GPCRs share the common structural organization consisting of an extracellular N-terminal, followed by seven transmembrane (7TM) α-helices that are connected via intra-and extracellular loops, and an unstructured intracellular C-terminal tail. GPCRs are designed to recognize and bind a variety of molecules thereby communicating changes in the extracellular environment with the interior of the cell. Together, these features have made them the most popular targets with 36% of all approved drugs targeting GPCRs^1,2^.

Upon agonist binding, GPCRs undergo a series of conformational changes that expose the GPCR intracellular cavity. The active receptor exhibits high affinity for heterotrimeric G proteins (Gαβγ), which bind to the GPCR intracellular cavity via the C-terminal α-helix of the Gα subunit (Fig. 1A)^3,4^. This event leads to G protein activation, marked by the exchange of GDP for GTP in the Gα subunit, dissociation of the Gα from the Gβγ subunits, and the initiation of downstream signaling pathways through effectors such as adenylyl cyclase and/or phospholipase C, leading to generation of secondary messengers including cyclic AMP (cAMP) and/or inositol 1,4,5-trisphosphate (IP_3_), respectively^5^. Ultimately, these signaling events affect cellular behavior and evoke physiological responses.

**Figure 1.**
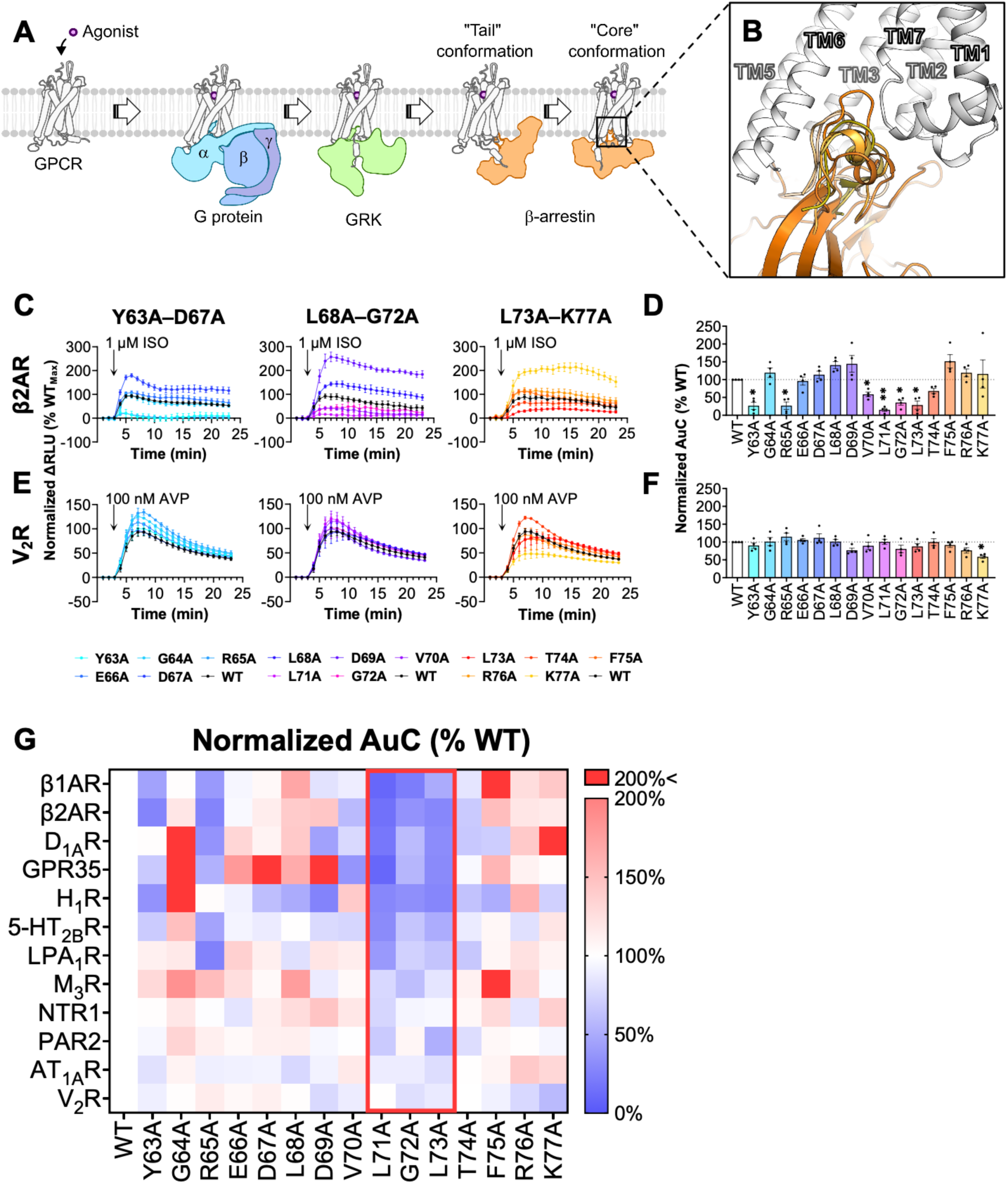
GPCR core interaction with the βarr1 finger loop. (**A**) Schematic illustration of associations between a GPCR and heterotrimeric G proteins, GRKs, and βarrs. βarr can engage with GPCRs in a “tail” and/or “core” conformations where the finger loop stabilize the core interaction. (**B**) Conformations adopted by residues on βarr1 finger loop within the GPCR cavity (GPCR backbone is adopted from PDB ID: 6TKO). (**C**) Changes in luminescence generated between βarr1-SmBiT finger loop mutants and β2AR-LgBiT in response to agonist stimulation (1 μM isoproterenol (ISO)). (**D**) The area under the curves (AuC) from the NanoBiT assay results are plotted as bar graphs. Data represents the means ± SEM of N=4 independent experiments, and one-way ANOVA with Dunnett’s multiple comparisons post hoc test was performed to determine statistical differences between βarr1-WT and its mutants (\**p* < 0.0332; \*\**p* < 0.0021). (**E**) Changes in luminescence generated between βarr1-SmBiT finger loop mutants and V_2_R-LgBiT in response to agonist stimulation (100 nM arginine vasopressin (AVP)). (**F**) The area under the curves (AuC) from the NanoBiT assay results are plotted as bar graphs. Data represents the means ± SEM of N=4 independent experiments, and one-way ANOVA with Dunnett’s multiple comparisons post hoc test was performed to determine statistical differences between βarr1-WT and its mutants (\**p* < 0.0332; \*\**p* < 0.0021). (**G**) Data from all GPCR-LgBiT/βarr1-SmBiT NanoBiT experiments are summarized in a heatmap highlighting increased (red) or decreased (blue) interaction between GPCRs and the βarr1 finger loop mutants.

Since prolonged or repeated agonist stimulation can be detrimental to the cells, they are equipped with elaborate mechanisms of GPCR desensitization. A general mechanism of desensitization involves GPCR kinases (GRKs) that recognize active receptor conformations and phosphorylate serines/threonines in the third intracellular loop (ICL3)^6^ and/or the C-terminal tail of the GPCR (Fig. 1A)^7,8^. Next, β-arrestins (βarrs) sense the phosphorylated receptor residues via lysine and arginine residues in their N-lobe, which leads to recruitment from the cytosol to the receptor at the plasma membrane (Fig. 1A)^9^. During GPCR–βarr complex formation, βarrs undergo conformational changes resulting in an extension of the finger loop (FL) between β-strands S5 and S6^10,11^. The extended FL can engage the GPCR intracellular cavity at an overlapping region to the G protein binding site (Fig. 1A)^12^. Thus, βarrs association with GPCRs in this configuration uncouples G proteins from the receptor, and thereby, terminates the G protein signaling^13–15^ (Fig. 1A). In addition to this, β-arrestins recruit components of the endocytic machinery such as AP-2 adaptors^16^ and clathrins^17^, which facilitate internalization of the GPCRs.

Over the past 20 years, advances in X-ray crystallography and cryo-electron microscopy (cryo-EM) have provided us with a near-atomic understanding of how GPCRs interact with its transducers and regulators. This is particularly true for their engagement with heterotrimeric G proteins where almost 1,000 high-resolution structures have been solved and deposited into the Protein Data Bank (PDB). On the other hand, the GPCR–βarr1 core engagement has only been visualized at high resolution for few GPCRs including the 5-hydroxytryptamine receptor 2B receptor (5-HT_2B_R), neurotensin receptor type 1 (NTR1), acetylcholine M2 receptor (M_2_R), cannabinoid receptor 1 (CB1R), vasopressin type 2 receptor (V_2_R), β1-adrenergic receptor (β1AR), μ-opioid receptor (μ-OR), apelin receptor (APJR), and GPR1^18–27^. Although these structures represent significant milestones in the field, the resolution in the βarr1-FL region is often limited due to its high flexibility, preventing precise modeling of the side chains involved in the interactions between the GPCR intracellular cavity and βarr1 (Fig. 1B). In addition, βarr1-FL residues that appear to participate in this core interaction have not been confirmed biochemically for most of these structures. Therefore, the exact manner by which the βarr1-FL interacts with the intracellular cavity of the diverse array of GPCR remain poorly understood. As this is unclear, it also remains a bit of a mystery how βarrs directly compete with G proteins for receptor binding considering that the βarr1-FL is an unstructured, flexible loop lacking any sequence similarity to the highly ordered C-terminal α-helix of the Gα subunits.

Using an interdisciplinary approach integrating systematic mutational mapping, bioinformatics, and structural analysis, we here sought to identify a universal binding mechanism by which the βarr-FL engages the intracellular GPCR core. Furthermore, we investigated whether this binding mechanism is shared by other transducers and regulators through an overlapping binding site within the GPCR core cavity. Such a mechanism may provide molecular and structural basis for how βarrs uncouple G proteins from the receptor, thereby contributing to the desensitization of G protein signaling.

## Results

### Biochemical characterization of βarr1-FL–GPCR interaction

Among the existing βarr isoforms, the FL consists of 15 well-conserved amino acids (Supplementary Fig. 1A). To enhance our general understanding of the βarr-FL engagement with the GPCR intracellular cavity, we made single alanine mutations of each individual residues of the βarr1-FL, expressed them equally in HEK293 cells, and investigated how βarr1-recruitment to a panel of 12 different GPCRs is affected using a split nanoluciferase (NanoBiT) approach (Supplementary Fig. 1B-C). Upon stimulation of each GPCRs with their respective agonists, a fast increase in the luminescence signal was observed for βarr1-WT indicating its recruitment to the GPCRs (Fig. 1C). Interestingly, different GPCRs exhibit varying sensitivity to mutations in the βarr1-FL (Fig. 1C-G and Supplementary Fig. 2). Additionally, the same βarr1-FL mutant can be effectively recruited to some receptors yet show poor recruitment to others (Fig. 1C-G and Supplementary Fig. 2). In general, GPCRs known to have a few sparse phosphorylation sites in their ICL3 and/or C-terminal tail (here referred to as Class A GPCRs) were more dramatically affected by the mutations, than those with cluster of phosphorylation sites (here referred to as Class B GPCRs) (Fig. 1G and Supplementary Fig. 2). Similar observations have been reported and attributed to the strong interactions between Class B GPCR phosphorylation clusters and βarrs, rendering the stability of these complexes less dependent on βarr-FL engagement with the GPCR intracellular cavity^28–30^. In addition, mutations at positions such as Y63 and R65 were frequently accompanied with reduced βarr1 recruitment (Fig. 1G and Supplementary Fig. 2). However, the residues that reduced βarr1 recruitment to GPCRs the most consistently were in a ‘hydrophobic tip’ composed of 71’-LGL-73’ (Fig. 1G and Supplementary Fig. 2). For 8 out of the 12 GPCRs tested, mutations in this hydrophobic tip significantly reduced recruitment to GPCRs suggesting that the dominating nature of the GPCR–βarr1 core interaction could be based on hydrophobic effect (Fig. 1G and Supplementary Fig. 2). As hydrophobic interactions are much less selective than polar interactions, it might also highlight how βarr1 couples universally to the wide range of GPCR cores, for which Gα subtypes are very selective. The 4 GPCRs minimally affected by the mutations in the hydrophobic tip are M_3_R, NTR1, angiotensin II type 1A receptor (AT_1A_R), and V_2_R. These receptors have Class B-type phosphorylation site clusters either in their ICL3 or C-terminal tail, and thus, are less dependent on the core engagement with βarrs to form stable GPCR–βarr complexes.

### Identification of a conserved leucine residues in GPCR-transducers interactions

To better understand the role of the hydrophobic FL tip (71’-LGL-73’), we reviewed all available experimentally solved structures of GPCR–βarr complexes in the core conformation. To date, such structural information is only available for 11 GPCRs including β_1_AR, NTR1, M_2_R, 5-HT_2B_R, μ-OR, GPR1, CB1R, PTHR1, rhodopsin, APJR and V_2_R. Interestingly, the βarr1 residues L71 and L73 are engaging the intracellular cavity in the majority of GPCRs (∼69%) at a hydrophobic patch defined by residues at TM3, TM5, and TM6 (Fig. 2A-C). We termed this configuration as *binding mode 1*. In case of CB1R, two structures are available, but βarr1-L71/L73 engage the receptor in binding mode 1 in only one of them (PDB ID 8WU1). In the other CB1R-βarr1 structure (PDB ID 8WRZ), only L73 engages with the hydrophobic patch, whereas L71 is pointed towards TM2 and TM7 (Fig. 2A and Supplementary Fig. 3A). We define this configuration as *binding mode 2*. Similar binding mode is observed in structures of rhodopsin-visual arrestin (bovine) complexes^31^. For the APJR, only L73 engages with the hydrophobic patch, whereas L71 is pointed outwards from the receptor center and is positioned in between TM6 and TM7/H8 (Fig. 2A and Supplementary Fig. 3D). We define this configuration as *binding mode 3*. Finally, the hydrophobic tip does not engage the receptor core profoundly in the V2R–βarr1 structure, which we define as *binding mode 4* (Fig. 2A and Supplementary Fig. 3G).

**Figure 2.**
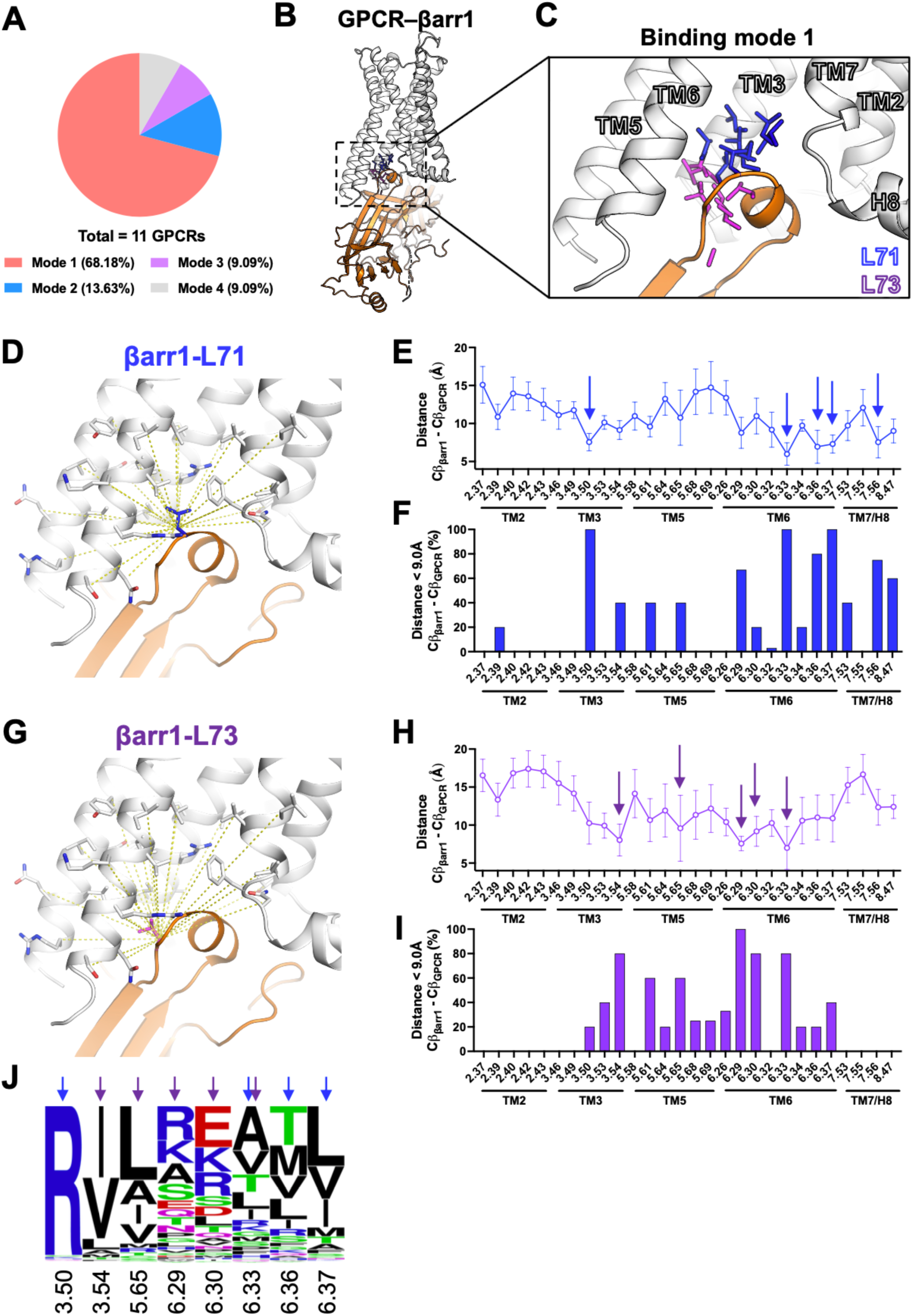
Structural analysis of the hydrophobic interaction between βarr1-L71 or βarr1-L73 and GPCRs. (**A**) Pie chart showing the percentage of βarr1-L71/L73 that engage with the GPCR intracellular cavity in different binding modes. (**B**) A representative GPCR–βarr1 structure in cartoon representation based on PDB ID: 7SRS, with (**C**) the relative positions of the βarr1-L71/73 residues in respect to the receptor in mode 1 from different structures highlighted in stick representations (PDB ID: 6TKO, 6UP7, 6U1N, 7SRS, 8WU1, 9LXR, 9UYH, and 9WSV). (**D**) Schematic for distance measurements between Cβ carbon of βarr1-L71 and Cβ carbons on all residues in the GPCR intracellular cavity. (**E**) Distance plot quantifying the average distances ± SD between Cβ carbon on βarr1-L71 and Cβ carbons of all residues within the GPCR intracellular cavity. (**F**) Bar graph showing the percentage of structures of which the distances between Cβ carbon of βarr1-L71 from each Cβ carbons on all residues in the GPCR intracellular cavity are below 9.0 Å. (**G**) Schematic for distance measurements between Cβ carbon of βarr1-L73 and Cβ carbons on all residues in the GPCR intracellular cavity. (**H**) Distance plot quantifying the average distances ± SD between Cβ carbon on βarr1-L73 and Cβ carbons of all residues within the GPCR intracellular cavity. (**I**) Bar graph showing the percentage of structures of which the distances between Cβ carbon of βarr1-L73 from each Cβ carbons on all residues in the GPCR intracellular cavity are below 9.0 Å. (**J**) Residue frequency plot showing the relative occurrence of residues in the GPCR intracellular cavity that interact with βarr1-L71/73. Residue positions closest to the βarr1-L71/L73 are highlighted with arrows in respective colors.

To determine likely receptor residues that partake in the hydrophobic association with the βarr1-FL hydrophobic tip, we measured distances between the Cβ carbons of βarr1-L71/L73 and all residues of the GPCR intracellular cavities (Fig. 2D-E and 2G-H and Supplementary Fig. 3B, 3E, and 3H). Additionally, we quantified the percentage of structures in which Cβ–Cβ distances between residues are below 9.0 Å, serving as an indicator of high likelihood of interaction (Fig. 2F and 2I and Supplementary Fig. 3C, 3F, and 3I). In binding mode 1, βarr1-L71 is positioned in proximity to H3.50, H6.33, H6.36, H6.37, and H7.56, whereas βarr1-L73 is closest to H3.54, H5.65, H6.29, H6.30, and H6.33 (Fig. 2D-J). In binding mode 2, βarr1-L71 is positioned in proximity to H2.39, H2.40, H7.56, and H8.47, whereas βarr1-L73 (methionine in visual arrestins) is closest to H3.50, H3.54, H6.32, H6.33, and H6.36 (Supplementary Fig. 3B-C). In binding mode 3, βarr1-L71 is positioned in proximity to H6.32, H6.33, H6.36, H7.56, and H8.47, whereas βarr1-L73 is closest to H3.54, H5.65, H5.68, H5.69, and H6.33 (Supplementary Fig. 3E-F). In binding mode 4, only the distance between βarr1-L71 and V2R-H2.37 is <9.0Å suggesting that the βarr1-FL does not engage robustly with the V_2_R intracellular cavity (Supplementary Fig. 3H-I).

**Figure 3.**
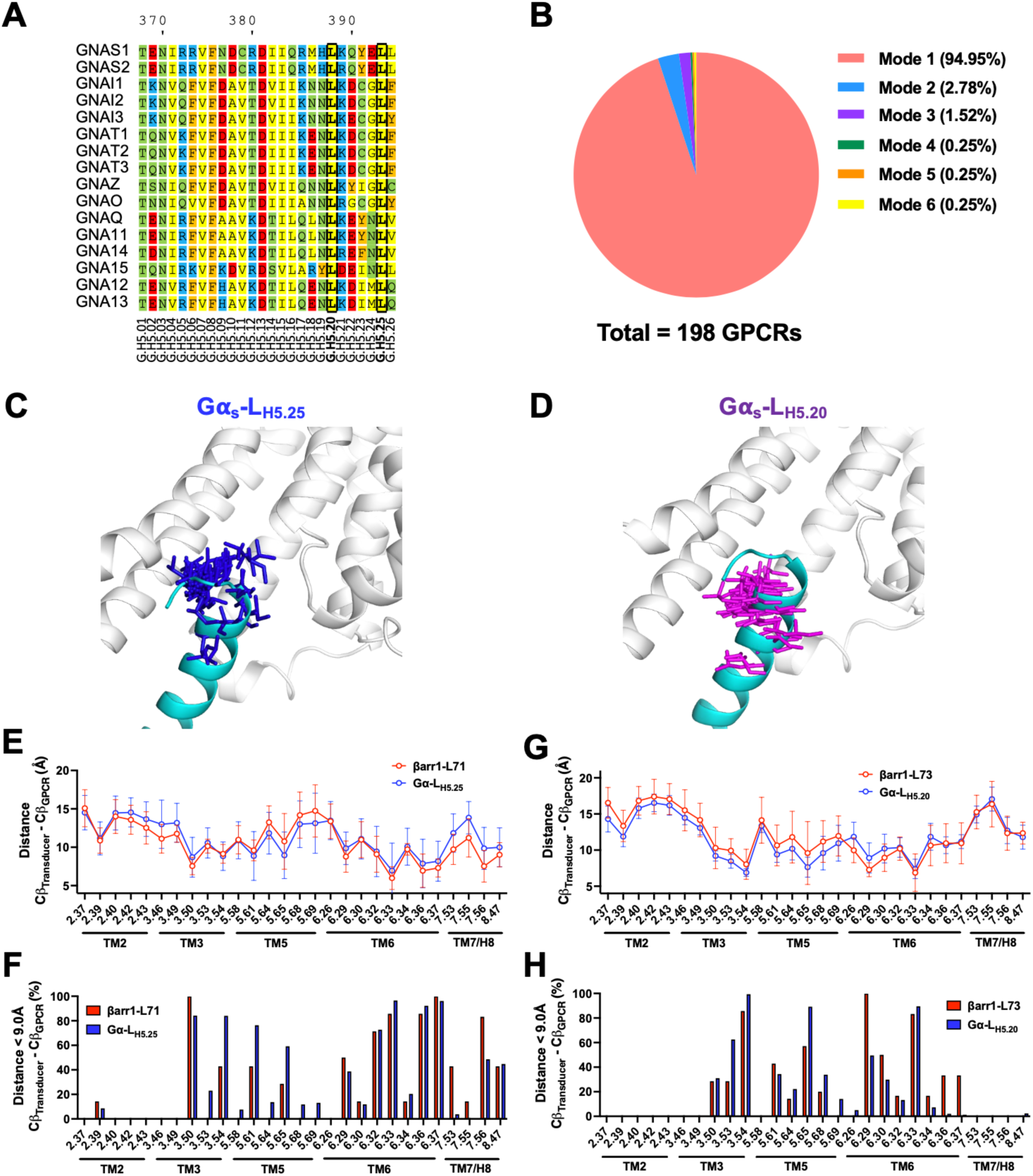
Comparison of the relative position within GPCR–transducer complexes between two highly conserved leucine residues on Gα subunits, H5.20 and H5.25, and βarr1-L71/L73. (**A**) Sequence alignment of the C-terminal helix H5 from all human Gα subtypes. (**B**) Pie chart showing the percentage of Gα subunit leucines (H5.20 and H5.25) that engage with the GPCR intracellular cavity in different binding modes. (**C** and **D**) Examples of GPCR–Gα_s_ protein structures superposed onto PDB ID: 3SN6 showing the relative positions of the leucine residues at H5.20 and H5.25 from different structures in respect to the receptor. (**E** and **G**) Distance plot quantifying the average distances ± SD between Cβ carbon of (**E**) Gα H5.25 or (**G**) Gα H5.20 and Cβ carbons of all residues within the GPCR intracellular cavity. Measured distances are compared with those between Cβ carbons of (**E**) βarr1-L71 or (**G**) βarr1-L71 and Cβ carbons of all residues within the GPCR intracellular cavity. (**F** and **H**) Bar graph showing the percentage of structures where the distances from Cβ carbon of (**F**) Gα H5.25 or (**H**) Gα H5.20 to each Cβ carbons on all residues within the GPCR intracellular cavity are below 9.0 Å. Percentages at each GPCR residues positions are compared with those from (**F**) βarr1-L71 or (**H**) βarr1-L71.

Interestingly, most of the structures of GPCR–G protein complexes (∼95%) show that the side chains of two strictly conserved leucine residues (L_H5.20_ and L_H5.25_) on the Gα C-terminal α-helix (H5) make similar *binding mode 1* hydrophobic contacts as βarr1-L71/L73 with hydrophobic patch in the GPCR intracellular cavity (Fig. 3A-D and Supplementary Fig. 4). Specifically, Gα residue L_H5.20_ is accommodated by GPCR residues H3.53, H3.54, H5.65, H6.29, and H6.33, while Gα residue L_H5.25_ is buried deeper in the receptor core facing the hydrophobic patch comprised by residues H3.50, H3.54, H6.33, H6,36, and H6.37 (Fig. 3C-H). The interaction mode of L_H5.20_ and L_H5.25_ with the GPCR intracellular cavity is very similar, if not identical, to that of the two βarr1-FL leucine residues (Fig. 3E-H). The strict conservation of these leucine residues on the C-terminal α-helix of the Gα subunit across all human paralogs suggests that they are essential for Gα function (Fig. 3A). Moreover, their universal role in receptor engagement indicates that they are unlikely to contribute to the Gα subtype selectivity exhibited by most GPCRs, a topic that has been extensively investigated by others^32–34^.

### Leucines H5.20 and H5.25 in the Gα C-terminal α-helix are critical for GPCR–G protein coupling

To investigate the role of the two strictly conserved Gα C-terminal α-helix leucine residues in GPCR–G protein coupling, we generated single and double alanine mutants whose recruitment to GPCRs were tested by the NanoBiT approach. As heterotrimeric G proteins naturally are present in the plasma membrane where the GPCRs are also already located, biophysical interactions between the two proteins are challenging to measure using cell-based proximity assays. To overcome this challenge, we used engineered miniG proteins instead, which has the distinct advantage of being expressed in the cytosol at resting states^35^. However, upon GPCR activation, these miniG proteins translocate to the receptors at the plasma membrane, which can be quantified effectively by the proximity assay (Fig. 4A). Most GPCRs we tested failed to recruit cognate miniG subtypes harboring alanine mutations of either L_H5.20_ (AL), L_H5.25_ (LA), or both (AA) upon agonist activation (expressed at comparable levels to the WT), with a curious exception of M_3_R, which significantly increased miniG protein recruitment (Fig. 4B-J and Supplementary Figs. 5 and 6).

**Figure 4.**
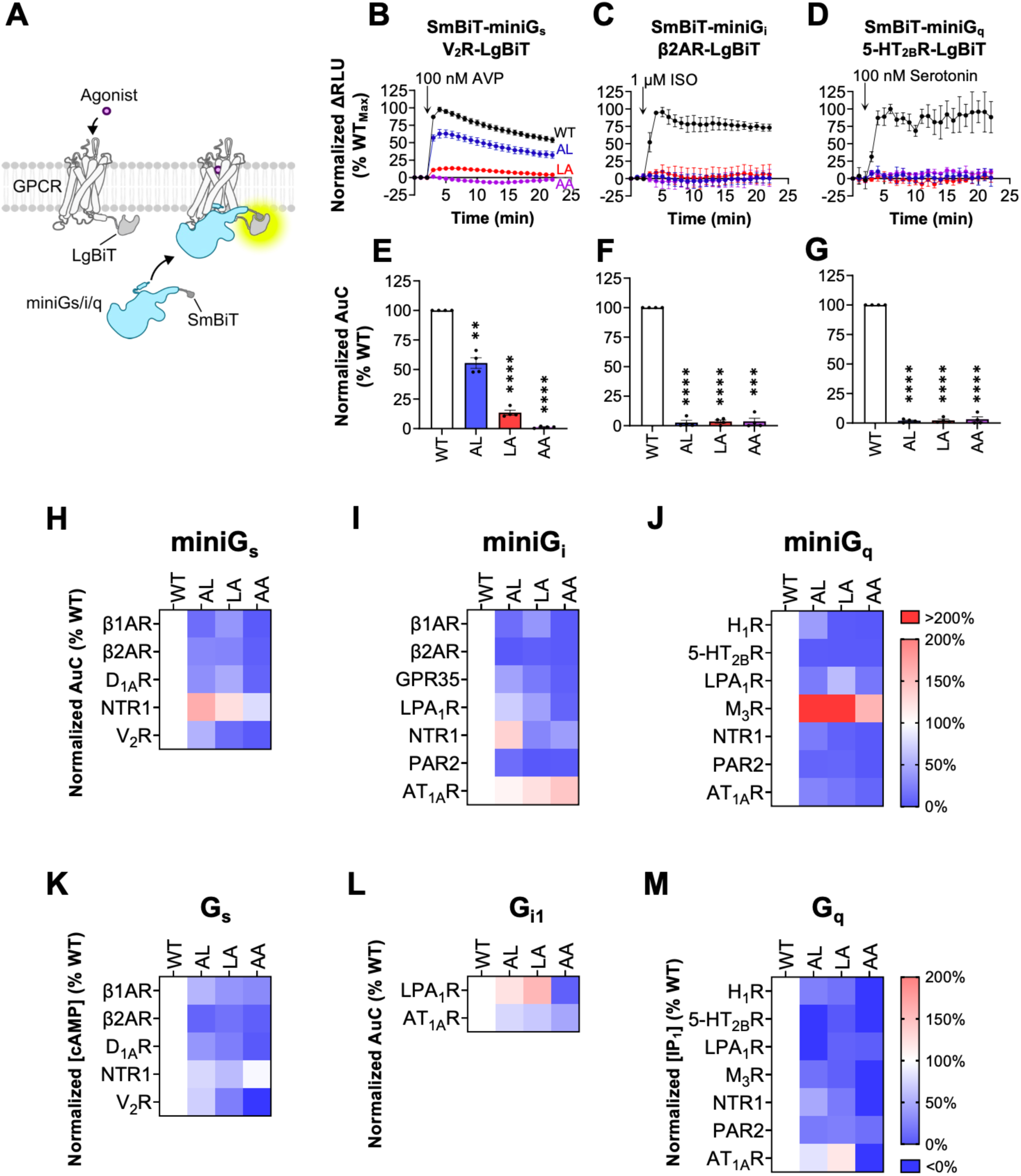
Importance of the Gα subunit C-terminal L_H5.20_ and L_H5.25_ residues for GPCR–G protein interaction and activation. (**A**) Schematic illustration of the NanoBiT assay used to measure interaction between GPCR-LgBiT and SmBiT-miniG protein mutants. (**B**-**D**) Changes in luminescence generated between (**B**) β2AR-LgBiT/SmBiT-miniG_s_, (**C**) β2AR-LgBiT/SmBiT-miniG_i_, or (**D**) 5-HT_2B_R-LgBiT/SmBiT-miniG_q_ in response to agonist stimulation (100 nM arginine vasopressin (AVP) for V_2_R, 10 μM isoproterenol (ISO) for β2AR, and 100 nM Serotonin for 5-HT_2B_R). (**E**-**G**) The area under the curves (AuC) from the corresponding NanoBiT assay results are plotted as bar graphs. Data represent the means ± SEM of N=4 independent experiments, and one-way ANOVA with Dunnett’s multiple comparisons post hoc test was performed to determine statistical differences between miniG-WT and miniG mutants (\**p* < 0.0332; \*\**p* < 0.0021;\*\*\**p* < 0.0002, \*\*\*\**p* < 0.0001). (**H**-**J**) Data from all GPCR-LgBiT and (**H**) SmBiT-miniG_s_, (**I**) SmBiT-miniG_i_, or (**J**) SmBiT-miniG_q_ NanoBiT experiments are summarized in heatmaps highlighting increased (red) or decreased (blue) interaction between GPCRs and the miniG protein mutants. (**K**-M) Data from GPCR-stimulated downstream signaling assays to detect (**K**) Gα_s_, (**L**) Gα_i1_, or (**M**) Gα_q_ activation is summarized in heatmaps highlighting increased (red) or decreased (blue) interaction between GPCRs and the Gα mutants.

In addition to measuring biophysical engagement between GPCRs and miniG proteins, we also verified potential decoupling events of the full-length heterotrimeric leucine mutants G proteins (expressed at similar levels to the WT) from the receptor by evaluating Gα-mediated downstream signaling events (Supplementary Fig. 7). Activity of Gα_s_ and Gα_q_ were assessed by measuring changes in intracellular levels of cAMP and inositol monophosphate (IP_1_, a stable metabolite of IP_3_ under the experimental conditions), respectively (Supplementary Fig. 8A-E and 8H-N). Signaling events through G_i/o_ was measured with the NanoBiT assay, by monitoring the recruitment of Rap1GAP to GPCRs in the presence of G_i1_ (Supplementary Fig. 8F-G). Rap1GAP only recognizes and binds to the activated G_i/o_ subtypes, and thus, this setup can be used to detect the ability of GPCRs to activate G_i/o_ proteins^36–38^. The downstream signaling assays showed almost identical outcome to the proximity assay, where the alanine mutants of the G proteins resulted in significantly reduced activity compared to the WT counterparts (even for M_3_R) despite being expressed at similar levels (Fig. 4K-M and Supplementary Fig. 8).

### Hydrophobic residues within the N-terminal α-helix of GRKs are critical for the GPCR-GRK interaction

Another important family of GPCR regulator is the GRKs. Upon receptor stimulation, GRKs senses the active conformation and gets recruited to the receptor where they phosphorylate the receptor ICL3 and/or C-terminal tail. In the case of GRK1–3, which are expressed in the cytosol, receptor stimulation leads to translocation of GRKs to the active receptor in the cell plasma membrane. On the other hand, GRK5/6 are naturally expressed in the plasma membrane and their recruitment to and phosphorylation of GPCRs are less dependent on receptor stimulation^39^. The GPCR–GRK interaction occurs via a conserved region on the GRK N-terminal α-helix that engages with GPCR intracellular cavity. To date, this interaction has only been visualized at high resolution in two cryo-EM structures of the rhodopsin–GRK1, and NTR1–GRK2–Gα_q_ complexes^40,41^, and thus, the molecular and ubiquitous nature of GPCR–GRK association is not well understood.

Comparisons between GPCR–G protein and GPCR–GRK structures reveal that the engagement of the GPCR core with Gα C-terminal and GRK N-terminal α-helices, respectively, overlaps to a great extent although the GPCR–GRK interaction is a bit more shallow (Fig. 5A)^40,41^. Given the ubiquitous nature of GRKs, we next focused on GRK residues that align with the hydrophobic patch in the GPCR intracellular cavity where the highly conserved leucine residues of βarrs-FL and Gα C-terminal α-helix bind. Interestingly, three aliphatic amino acid residues on the GRKs N-terminal α-helix face the hydrophobic patch in a similar manner to the leucines (Fig. 5B-D). The hydrophobic GRK residues are not as strictly conserved as seen in the other transducers except for L_H1.02_, but leucines, isoleucines, or valines at positions H1.05 and H1.06 sufficiently compose the hydrophobic interface (Fig 5B). Residues of the hydrophobic patch on the receptor that seem to comprise the interface for GRK engagement include H3.54, H5.65, H5.68, H6.26, H6.29, H6.30, H6.32, H6.33, H6.36, H7.56, and H8.47 (Fig. 5C-D).

**Figure 5.**
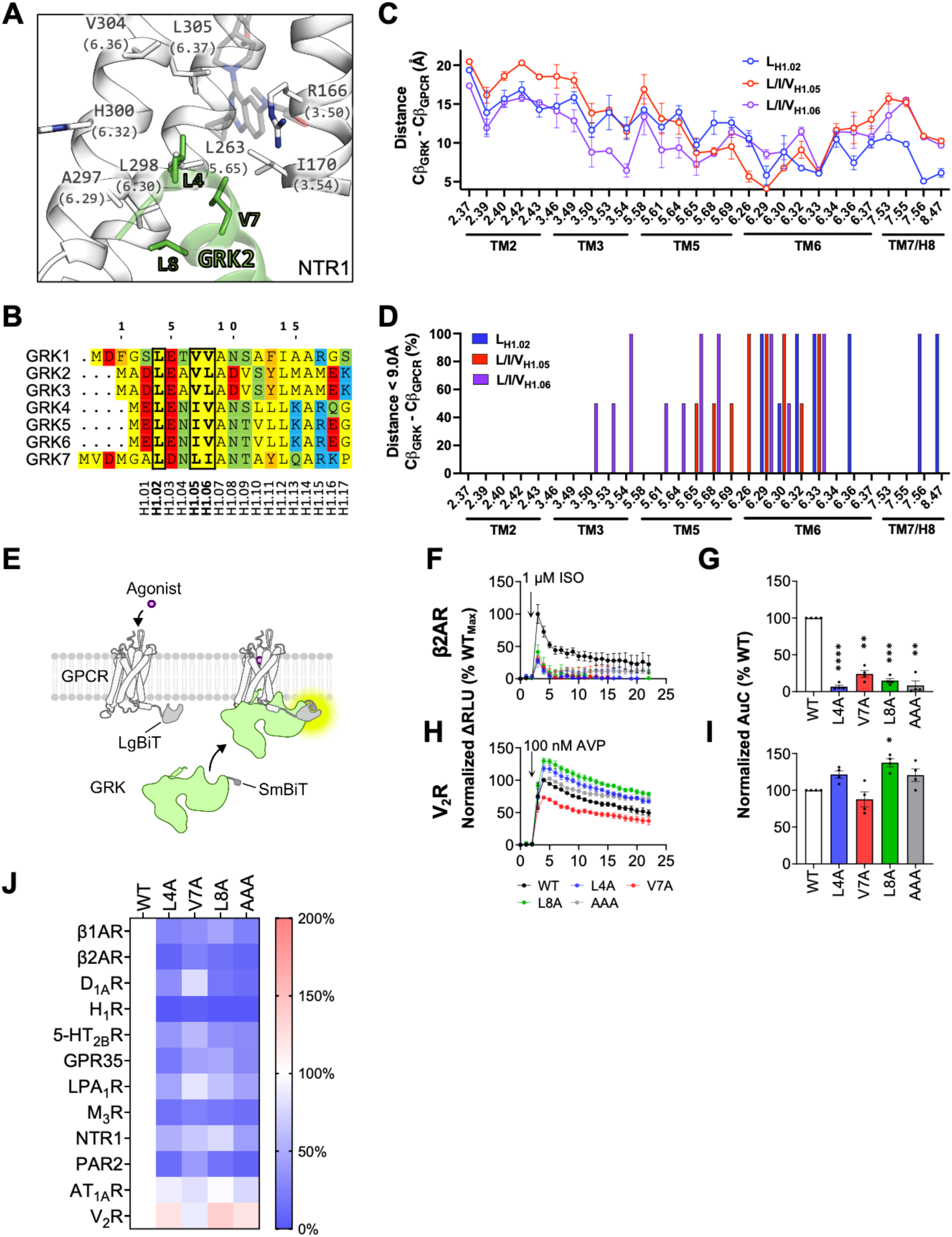
Dependency of the GRK N-terminal aliphatic residues at position H1.02, H1.05, and H1.06 for the GPCR–GRK interaction. (**A**) Structural model displaying the interaction of N-terminal α-helix of GRK2 with the intracellular NTR1 cavity (adopted from PDB ID: 8JPB). Generic receptor residue numbers (Ballesteros-Weinstein numbering scheme) are shown in brackets. (**B**) Multiple sequence alignment of human GRK paralogs. GRK2 residue numbering (top) and generic residue numbering (bottom) are shown. Aliphatic residues at positions H1.02, H1.05, and H1.06, which are involved in the GPCR engagement, are highlighted in bold. (**C**) Distance plot quantifying the average distances ± SD between Cβ carbons of GRK1/2-L_H1.02_/V_H1.05_/L/V_H1.06_ and Cβ carbons of all residues within the GPCR intracellular cavity. (**D**) Bar graph showing the percentage of structures where the distance between Cβ carbon of GRK1/2-L_H1.02_/V_H1.05_/L/V_H1.06_ and each individual Cβ carbons of all residues within the GPCR intracellular cavity is below 9.0 Å. (**E**) Schematic illustration of the NanoBiT assay applied to measure interaction between GPCR-LgBiT and GRK2-SmBiT mutants. (**F** and **H**) Changes in luminescence generated between GRK2-SmBiT N-terminal mutants and (**F**) β2AR-LgBiT or (**H**) V_2_R-LgBiT in response to agonist stimulation (1 μM isoproterenol (ISO) for β2AR and 100 nM arginine vasopressin (AVP) for V_2_R). (**G** and **I**) The area under the curves (AuC) from the NanoBiT assay results are plotted as bar graphs. Data represent the means ± SEM of N=4 independent experiments, and one-way ANOVA with Dunnett’s multiple comparisons post hoc test was performed to determine statistical differences between GRK2-WT and mutants (\**p* < 0.0332; \*\**p* < 0.0021; \*\*\**p* < 0.0002; \*\*\*\**p* < 0.0001). (**E**) Data from all GPCR-LgBiT/GRK2-SmBiT NanoBiT experiments are summarized in a heatmap highlighting increased (red) or decreased (blue) interaction between GPCRs and the GRK2 N-terminal mutants.

**Figure 6.**
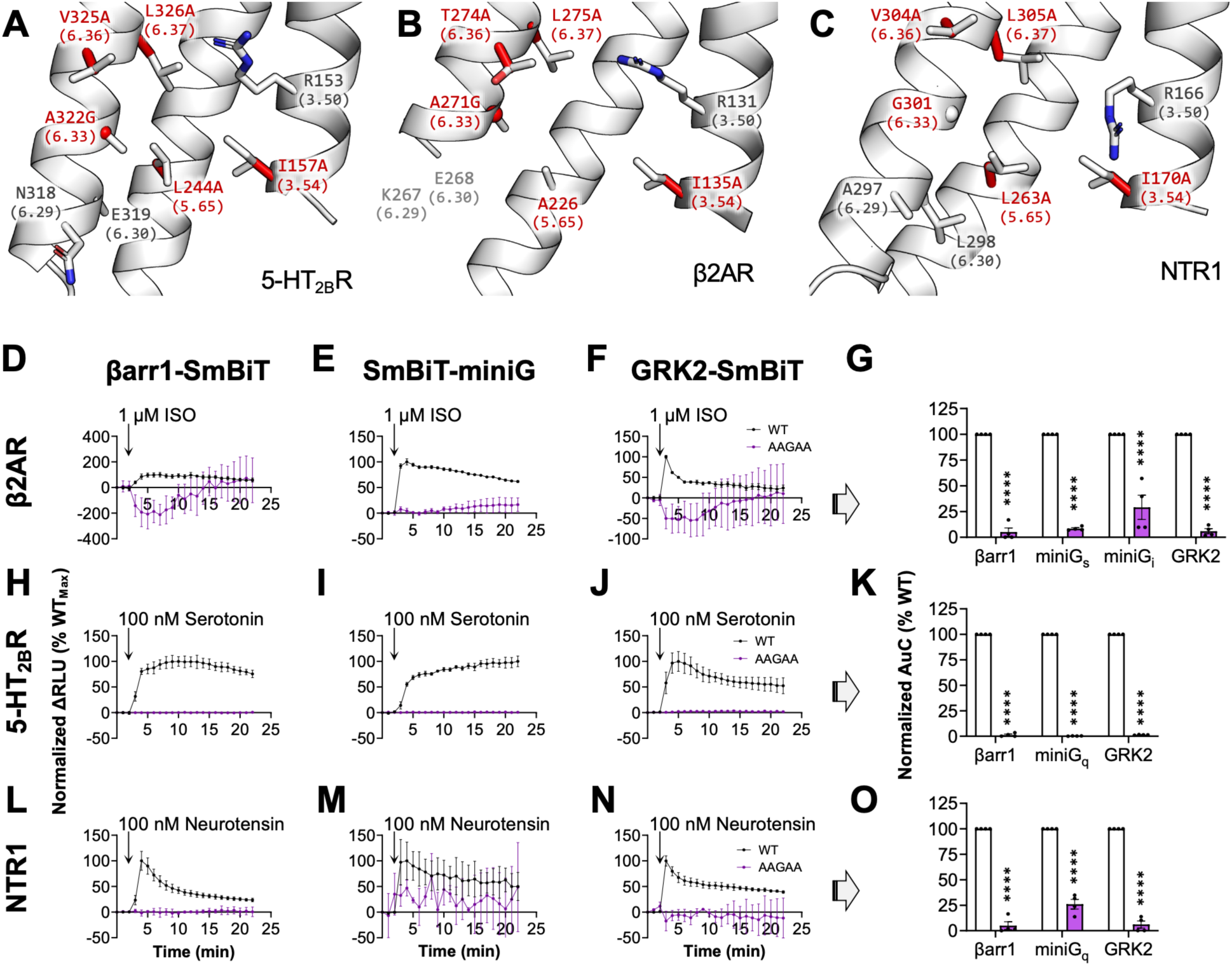
A common hydrophobic path within the GPCR intracellular cavity engage with the conserved transducer/GRK aliphatic residues. (**A**) Structural models displaying a common hydrophobic patch in the GPCR intracellular cavity made up of aliphatic residues shown at positions H3.54, H5.65, H6.33, H6.36, and H6.37 (adopted from PDB ID: 7SRS, 8GG0, and 8JPB). Generic receptor residue numbers (Ballesteros-Weinstein numbering scheme) are shown in brackets. Stick representations in red show the amino acid residues they were mutated into for generation of the AAGAA mutants. (**D–F**) 1 μM isoproterenol (ISO)-stimulated changes in luminescence generated between WT or AAGAA mutant β2AR-LgBiT and (**D**) βarr1-SmBiT, (**E**) SmBiT-miniGs, or (**F**) GRK2-SmBiT. (**H–J**) 100 nM serotonin-stimulated changes in luminescence generated between WT or AAGAA mutant 5-HT_2B_R-LgBiT and (**H**) βarr1-SmBiT, (**I**) SmBiT-miniGq, or (**J**) GRK2-SmBiT. (**L–M**) 100 nM neurotensin-stimulated changes in luminescence generated between WT or AAGAA mutant NTR1-LgBiT and (**L**) βarr1-SmBiT, (**M**) SmBiT-miniGq, or (**N**) GRK2-SmBiT. (**G, K** and **O**) The area under the curves (AuC) from the NanoBiT results from the (**G**) β2AR, (**K**) 5-HT_2B_R, and (**O**) NTR1 experiments are plotted as bar graphs. Data represent the means ± SEM of N=4 independent experiments, and students *t* tests were performed to determine statistical differences between WT and AAGAA GPCRs across all transducers (\*\*\*\**p* < 0.0001).

To investigate the general importance of the conserved aliphatic residues within GRK2/3 N-terminal α-helix, we mutated them individually to alanine and expressed WT and the mutant GRK2-SmBiT constructs to a similar extent in HEK293 cells (the N-terminal α-helices of GRK2 and GRK3 are identical) (Supplementary Fig. 9). In addition, GPCR-LgBiT constructs were co-expressed and agonist-stimulated GRK2 recruitment was measured using NanoBiT approach (Fig 5E). In accordance with the results from the other transducers, mutating individual or all three aliphatic residues of the GRK2 N-terminal α-helix that face the GPCR hydrophobic patch resulted in significant decrease in receptor engagement across most of the 12 GPCRs tested against (Fig. 5F-J and Supplementary Fig. 10). Interestingly, both AT_1A_R and V_2_R was minimally affected by the GRK2 mutations, resembling what was observed from the βarr1 mutants (Fig. 5H-J and Supplementary Fig. 10K-L).

### Identified key hydrophobic residues on βarr1, Gα subunit, and GRKs engage a conserved hydrophobic patch in the intracellular cavity of active GPCRs

Sequence alignment of all human GPCRs revealed that the hydrophobic patch in the receptor core is highly conserved in terms of its physiochemical nature (Fig. 2J). Our mutagenesis experiments and structural analyses of GPCR–transducer complexes suggest that common transducer motifs bind the hydrophobic patch in the GPCR intracellular cavity (Figs. 1-5), and thus, directly provide a molecular basis for the competition between the transducers that leads to desensitization of G protein signaling.

To further substantiate a central role for this hydrophobic patch in GPCR interaction with transducers/regulators, we sought to reduce the capacity for hydrophobic contacts within the intracellular receptor cavity of three representative GPCRs: β2AR, 5-HT_2B_R, and NTR1. Five residues that line the GPCR–transducer/regulator interface and are mostly hydrophobic in all three receptors, H3.54, H5.65, H6.33, H6.36, and H6.37, were selected for alanine substitution (Fig. 6A-C and Fig. 3E-H). If the natural amino acid at position H6.33 is already an alanine, the residue was mutated to glycine. Computational introduction of these mutations into the corresponding GPCR-transducer/regulator structures, either experimentally solved or predicted by AlphaFold3 (when experimental structures were unavailable), resulted in a substantial decrease in the buried surface area (BSA) of interacting leucine/aliphatic residues on Gα, βarr1, and GRK2 (Supplementary Fig. 11). This observation indicates that the mutations disrupt the hydrophobic association between the intracellular GPCR hydrophobic patch and transducers/regulators.

To test this hypothesis, the surface expression levels of the wild-type and mutant GPCRs in HEK293 cells were first matched as determined by ELISA (Supplementary Fig. 12), followed by NanoBiT measurement of agonist stimulated recruitment of miniG protein(s), βarr1, and GRK2. As expected, the effect of the receptor quintuple mutant (AAGAA) on the transducer recruitment was detrimental (Fig. 6D-O). All in all, these results suggest that the interface composed of the conserved hydrophobic residues on the transducers/regulators of diverse sequences and the hydrophobic patch in the intracellular GPCR cavity provide molecular basis for their nondiscriminatory interactions.

## Discussion

Upon agonist stimulation, GPCRs undergo a conformational change that exposes the GPCR intracellular cavity. The GPCR cavity is the site of interaction common for all three families of its transducers/regulators. Gα subunits and GRKs utilize their C-terminal and N-terminal helices, respectively, while βarrs extend their FL for the engagement to the GPCR cavity. Despite substantial differences in the sequences and secondary structures of the receptor-interacting regions of the transducers, they bind to and compete for the exposed GPCR cavity. This competition for receptor engagement between G protein and βarrs in particular is essential for desensitization of G protein signaling as βarrs uncouple G proteins from the active receptors that has been phosphorylated^13,14^. For decades, it has been hypothesized that there exists an overlapping binding site within the intracellular GPCR cavity where a direct competition between G proteins and βarrs takes place. However, the identity of such overlapping binding site has not been generally described prior to this study. Using a combination of systemic mutational mapping, bioinformatics, and extensive structural analysis, we accumulated supporting evidence for such overlapping binding sites consisting of a hydrophobic patch formed in general by residues located on TM3 (H3.54), TM5 (H5.65), and TM6 (H6.29, H6.30, H6.33, H6.36, and H6.37) of GPCRs. A previous biochemical study also identified residues (L226 (H5.61) and V230 (H5.65)) at/close to this hydrophobic patch in bovine rhodopsin as a hot spot for engagement with Gα_t_ subunit, visual arrestin, and GRK1. In addition, changing the aliphatic nature of H3.5l4, H5.65, H6.33, and H6.37 in β2AR to hydrophilic via mutation has been found to negatively affects the receptor’s ability to activate G_s_ protein^42^. Expanding on these studies, we showed that mutating key residues of the hydrophobic patch (H3.54, H5.65, H6.33, H6.36, and H6.37) within the human β2AR, 5-HT_2B_R, and NTR1 also led to a dramatic reduction in the ability of the receptors to recruit G protein, βarr1, and GRK2 (Fig. 6).

Another major objective of the study was to elucidate the general features of the βarr1-FL interaction with GPCRs. Unlike GPCR–G protein complexes, structural information on GPCR–βarr complexes in the’core’ conformation remains limited, largely due to technical challenges in stabilizing these complexes in this state. Moreover, in the few structures that have been solved, the resolution of the βarr1-FL region varies significantly owing to the intrinsic flexibility of the FL^18,19^. As a result, the full side chains of FL residues have been modeled in only a subset of these complex structures^19^, leaving high-resolution insights into the βarr1-FL core interaction still largely lacking. To identify key residues within βarr1-FL that partake in the GPCR–βarr1 core interaction, we conducted alanine-scanning mutagenesis and demonstrated that the hydrophobic tip consisting of 71’-LGL-73’ plays an important general role (Fig. 1). For most GPCRs tested, we found that a set of mutations in other FL residues also influenced βarr1 recruitment negatively and in a few cases positively (Fig. 1C-G). However, the specific pattern of residues involved were, in most cases, unique to each GPCR. Similar findings has been made in the rhodopsin–visual arrestin system where mutation of residues in the hydrophobic tip of visual arrestin (75’-MGL-77’ in bovine visual arrestin), as well as other residues significantly reduced binding to the activated receptor^12,43^.

By unbiased Cβ–Cβ distance measurement between βarr1 L71 or L73 and all residues comprising the intracellular GPCR cavities, we found these FL leucines are in proximity to residues of the hydrophobic patch on the receptor in most of available GPCR–βarr1 structures (binding mode 1, Fig. 2). Interestingly, in 95% of the GPCR–G protein structures evaluated, two conserved leucine residues on helix H5 at the C-terminal of all Gα subtypes (H5.20 and H5.25) appear to engage with the receptor hydrophobic patch in a similar, if not identical manner (Fig. 3). These residues are highly conserved among all Gα subtypes (Fig. 3A) and across different species suggesting that they play an essential role in the coupling to GPCRs^32,44^. The critical role of these leucines in Gα_i_ for interacting with rhodopsin has been documented previously^45^, but our findings demonstrate their universal importance as ‘interaction hotspots’ not only in the recruitment of major G protein subtypes to active GPCRs, but also the GPCR–βarr core interaction (Figs. 1-4). Interestingly, inspection of structures of the only two GPCR–GRK complexes available revealed that 3 aliphatic residues, L_H1.02_, V_H1.05_, and L/V_H1.06_ of the GRK1/2 N-terminal α-helix also associate with the receptor hydrophobic patch, although the relative positions of these aliphatic side chains with respect to the GPCR intracellular cavity differ somewhat as compared to the conserved leucine residues in both the βarrs and Gα subunits (Fig 5A-D). Importantly, mutation of these residues to alanine in GRK2 significantly reduced their recruitment to most GPCRs tested suggesting that a hydrophobic association between the GRK N-terminal and the receptor hydrophobic patch is essential for the stability of the complex (Fig. 5E-J). Reduction of GRK2’s ability to phosphorylate active rhodopsin has also previously been linked to mutation of these residues further substantiating the importance of this interaction^46^.

Surprisingly, the recruitment of GRK2 to two Class B GPCRs, AT_1A_R and V_2_R, was insensitive to mutations in the L_H1.02_, V_H1.05_, and/or L_H1.06_ (Fig. 5J and Supplementary Fig. 10K-L). Although the reason for this insensitivity was not further explored in this study, it has long been known that free Gβγ generated, as a result of G protein activation, plays a key role in the GRK2/3 recruitment to and phosphorylation of GPCRs^47,48^. We and others have shown that Gβγ subunits bind to βarrs when in complex with active V_2_R or PTHR1 to form a stable GPCR–βarr–Gβγ “miniplex”^49–51^. Such stable miniplexes could act as scaffold for which the recruitment of GRK2/3 via the Gβγ subunits could take place. In this scenario, the GRK2/3 N-terminal α-helix interaction with the GPCR intracellular cavity would be less important for the overall GRK2/3 recruitment.

Over the past 10-15 years, it has become increasingly evident that βarrs can engage with GPCRs in multiple distinct conformations. One such binding mode is the ‘tail’ conformation, in which βarrs only interact with heavily phosphorylated clusters on the C-terminal tail or ICL3 of Class B GPCRs^28–30^. Interestingly, in this conformation, βarrs still maintain their ability to facilitate key functions such as receptor endocytosis and βarr-mediated signaling^28–30^, but not desensitization of G protein signaling^28^. To our surprise, a recent structural study of the atypic chemokine receptor in complex with βarr1 revealed an even more unconventional binding mode^52^. In this structure, the βarr1-FL does not engage the hydrophobic patch in the GPCR core. Instead, it inserts into micellar boundary near TM1 and TM7 of the receptor. Although a previous study reported FL insertion into the plasma membrane to stabilize an active and receptor-free state of βarr association with the membrane^53^, it was unexpected to see this FL insertion into the membrane even when βarr1 is in complex with a GPCR. Curiously, these new findings indicate that the hydrophobic tip of the βarr-FL may be so promiscuous that it fails to distinguish not only between different GPCR cores, but also between the receptor core and the surrounding lipid membrane environment.

## Methods

### Reagents

3-isobutyl-1-methylxanthine (IBMX; cat. no. I5879), angiotensin II (cat. no. A9525), bovine serum albumin (BoSA; cat. no. A2153), carbachol (cat. no. C4382), dopamine (cat. no. H8502), forskolin (cat. no. F6886), histamine (cat. no. 53300), isoproterenol (ISO; cat. no. 42035), mouse anti-FLAG M2-HRP antibody (cat. no. A8592), neurotensin (cat. no. N6383), oleoyl-L-α-lysophosphatidic acid (LPA; cat. no. L7260), paraformaldehyde (PFA; cat. no. 158127), serotonin (cat. no. H9523), trypsin (cat. no. T0303), Tween-20 (cat. no. P1379), and zaprinast (cat. no. Z0878) were all purchased from Sigma-Aldrich. Dulbecco’s Modified Eagle Medium (DMEM) with GlutaMAX™ supplement (cat. no. 10566016), fetal bovine serum (FBS; cat. no. A5670801), Hanks’ Balanced Salt Solution (HBSS; cat. no. 14025076), Lipofectamine 3000 transfection reagent (cat. no. L3000-015), Opti-MEM (cat. no. 31985070), parental HEK293 cell line (cat. no. R70507), penicillin-streptomycin (cat. no. 15140122), SuperSignal West Pico PLUS chemiluminescent substrate (cat. no. 34580), and trypsin–EDTA 0.05% (cat. no. 25300054) were obtained from Thermo Fisher Scientific. Arg8-vasopressin (AVP; cat. no. ab120175) was obtained from Abcam. 96-well white flat-bottom plates (cat. no. 353296) were purchased from Corning. 384-well small-volume white flat-bottom plates (cat. no. 784075) were from Greiner Bio-One. Coelenterazine-h (301) was obtained from NanoLight. Skim milk powder (cat. no. M17200) was purchased from RPI Research Products. Mouse anti-LgBiT antibody (cat. no. N7100) and rat anti-SmBiT antibody were obtained from Promega. Sheep anti-mouse IgG HRP-linked F(ab’)₂ (cat. no. NA9310V) and donkey anti-rabbit IgG HRP-linked F(ab’)₂ (cat. no. NA9340V) were purchased from Cytiva (Amersham). Goat anti-rat IgG-HRP (cat. no. HAF005) was obtained from R&D Systems. Mouse anti-Gα_s_ antibody (cat. no. sc-135914) was purchased from Santa Cruz Biotechnology. Rabbit anti-Gα_i_ (cat. no. PA5-28223) and rabbit anti-Gα_q_ (cat. no. PA5-17693) antibodies were obtained from Invitrogen. HTRF cAMP detection reagents, including cAMP Eu-cryptate antibody and cAMP-d2 (cat. no. 62AM4PEB), and IP-One detection reagents, including IP1 Tb-cryptate antibody and IP1-d2 (cat. no. 62IPAPEB), were purchased from Revvity. The βarr1/2 double knockout and total Gα knock-out derivatives were gifts from Dr. Asuka Inoue (Tohoku University). Rabbit anti-β-arrestin 1 antibody (A1CT) was a gift from Dr. Robert J. Lefkowitz (Duke University, Durham, North Carolina).

### Plasmid constructs

Human GPCRs (β1AR, β2AR, D_1A_R, GPR35, H_1_R, 5-HT_2B_R, LPA_1_R, M_3_R, NTR1, PAR2, AT_1A_R, and V_2_R) N-terminally tagged with an influenza hemagglutinin signal sequence (MKTIIALSYIFCLVFA) followed by a FLAG-tag (DYKDDDDK) and C-terminally tagged with an LgBiT through a flexible linker (GGSGGGGSGGSSSGG) were synthesized and cloned into pTwist CMV vectors by Twist Bioscience. Human β2AR, 5-HT_2B_R, and NTR1 quintuple mutants (AAGAA) tagged as above were synthesized and cloned into pTwist CMV vectors by Twist Bioscience. Human βarr1 N-terminally tagged with a SmBiT through a flexible linker (GGSGGGGSGGSSSGG) and its finger loop mutant derivatives were synthesized and cloned into pTwist CMV vectors by Twist Bioscience. MiniGα proteins N-terminally tagged with a nuclear export signal (MLQNELALKLAGLDINKT) and an influenza hemagglutinin (HA)-tag (YPYDVPDYA) followed by a SmBiT each connected with flexible linkers (GGSG) were synthesized and cloned into pTwist CMV vectors by Twist Bioscience. Human Gα proteins were synthesized and cloned into into pTwist CMV vectors by Twist Bioscience. Human GRK2 C-terminally tagged with a SmBiT through a flexible linker (GGSGGGGSGGSSSGG) were synthesized and cloned into pTwist CMV vector by Twist Bioscience. Alanine mutants of miniGα, Gα, and GRK were generated using the Q5® Site-Directed Mutagenesis Kit with mutagenesis primers synthesized by IDT.

### Cell cultures and Transfection

Parental HEK293 cell line and its knock-out derivatives (βarr1/2 double K.O. and total Gα K.O. cell lines) were cultured in DMEM GlutaMAX™ supplemented with 10% (v/v) FBS and 100 U/ml Penicillin-Streptomycin at 37 °C and 5% CO₂. Cell cultures were passaged every 3–4 days using trypsin–EDTA 0.05%. For experiments, cultures with up to 15 passages were used. Transfections of DNA constructs were performed according to the manufacturer’s protocol.

### NanoBiT assay

NanoBiT assays between GPCR and transducer pairs each tagged with LgBiT and SmBiT, respectively, were performed to measure the interaction between the proteins. HEK293 cells transfected with GPCR-LgBiT and transducer-SmBiT were plated on poly-D-lysine coated White Flat Bottom Microplate and equilibrated in Opti-MEM™ at 37°C for 60 minutes. Coelenterazine-h was added at a final concentration of 10 μM before starting the measurement. After establishing a baseline response for 2 minutes, cells were stimulated with respective agonists added at appropriate concentrations and the luminescence was measured for another 20 minutes. 10 μM ISO for β1AR and β2AR, 100 nM dopamine for D_1A_R, 10 μM Zaprinast for GPR35, 100 nM histamine for H_1_R, 100 nM serotonin for 5-HT_2B_R, 10 μM LPA for LPAR1, 10 μM carbachol for M_3_R, 100 nM neurotensin for NTR1, 100 nM trypsin for PAR2, 1 μM angiotensin for AT_1A_R, and 100 nM AVP for V_2_R were used to stimulate the receptors. The signal was detected at 550 nm using a PHERAstar *FSX* instrument (BMG LabTech). No blinding from the different conditions was done. NanoBiT assays between GPCRs-LgBiT and Rap1GAP-SmBiT with G_i1_ (wild-type or alanine mutants) were performed as above to measure G_i_ activation from GPCR stimulation with respective agonists. NanoBiT assays between βarr1-LgBiT and SmBiT-miniG_q_ (wild-type or alanine mutants) were performed as above to measure miniG_q_ expression in HEK293 total Gα K.O. cell line.

### HTRF cAMP Gs Assay

HEK293 total Gα K.O. cells transfected with GPCRs and G_s_ (wild-type or alanine mutants) were harvested and resuspended in assay buffer (20 mM HEPES in HBSS buffer, pH 7.4 supplemented with 500 nM IBMX). Ligand buffer (assay buffer supplemented with 20 μM forskolin) containing agonists at appropriate concentrations, or vehicle, was mixed with the cell suspension in 384-well plates. The plates were incubated at 37°C for 30 minutes before adding lysis buffer containing cAMP Eu-cryptate antibody working solution and cAMP-d2 reagent working solution, followed by an additional incubation for 1 hour at room temperature. The fluorescence was measured on the PHERAstar *FSX* where wells were excited with light at 340 nm and emission light was measured at 615 nm and 665 nm. The TR-FRET 665 nm/615 nm ratio, which is inversely proportional to the cAMP concentration, was used in combination with a cAMP standard curve to calculate the cAMP production in the cells.

### HTRF IP-One Gq Assay

HEK293 total Gα K.O. cells transfected with GPCRs and G_q_ (wild-type or alanine mutants) were harvested and resuspended in assay buffer. StimB buffer containing agonists at appropriate concentrations, or vehicle, was mixed with cell suspension in 384-well plates. The plates were incubated at 37°C for 30 minutes before adding lysis buffer containing IP1 Tb-cryptate antibody working solution and anti IP1-d2 reagent working solution, followed by an additional incubation for 1 hour at room temperature. The fluorescence was measured on the PHERAstar *FSX* where wells were excited with light at 340 nm and emission light was measured at 615 nm and 665 nm. The TR-FRET 665 nm/615 nm ratio, which is inversely proportional to the IP_1_ concentration, was used in combination with a IP_1_ standard curve to calculate the IP_1_ production in the cells.

### Western blotting

Western blotting was conducted to confirm correct expression of GPCRs, transducers, and their mutants. All blots were probed with the primary antibody diluted 1:1000 in blocking buffer (5% dry milk powder in 1×TBS supplemented with 1% (v/v) Tween**^®^** 20) for 16 hours at 4°C followed by incubation with HRP-conjugated secondary antibodies diluted at 1:1000 in the blocking buffer. Blots visualized using SuperSignal^™^ West Pico PLUS Chemiluminescent Substrate on a ChemiDoc™ XRS+ Imaging System (Bio-Rad).

### ELISA cell surface expression

Cell surface expression of wild-type and the quintuple mutants of β2AR, NTR1, and 5-HT_2B_R were evaluated using ELISA. HEK293 cells were seeded on poly-D-lysine coated 96-well plates and incubated overnight. Cells were fixed with 4% PFA and blocked with 2% (w/v) BoSA in PBS, before incubating 1 hour at room temperature with mouse anti-FLAG M2-HRP antibody diluted 1:10,000 in blocking buffer. Enzymatic chemiluminescence was generated by the addition of SuperSignal™ West Pico PLUS as substrate and measured on a CLARIOstar plate reader (BMG LabTech) with the emission filter 410-80.

### Structural analyses

Structures of the high resolutions were retrieved from the Protein Data Bank for analysis (see all models used for this study in ‘Binding modes.xlsx’). The assignment of binding modes of the conserved leucine residues at position 71 and 73 on βarr1 (and visual arrestin) and H5.20 and H5.25 on Gα was classified based on the overall position relative to the TM α-helices as described. A small number of GPCR structures exhibit two alternative binding modes for the transducer leucine residues. For these structures, each binding mode was assigned a weight of 0.5 for statistical purposes. A more precise determination of the positions of these residues, as well as GRK residues H1.02, H1.05, and H1.06, relative to residues forming the GPCR intracellular cavity was achieved through distance measurements. To this end, distances between the Cβ carbons on the transducer leucine residues (L71 and L73 on arrestins and H5.20 and H5.25 on Gα) or hydrophobic residues of GRKs and the Cβ carbons of every residue within the corresponding GPCR intracellular cavity (2.37, 239, 2.40, 2.42, 2.43, 3.46, 3.49, 3.50, 3.53, 3.54, 5.58, 5.61, 5.64, 5.65, 5.68, 5.69, 6.26, 6.29, 6.30, 6.32, 6.33, 6.34, 6.36, 6.37, 7.53, 7.55, 7.56, and 8.47) were measured in the PyMOL Molecular Graphics System (Version 3.0 Schrödinger, LLC). The GPCRs were assigned with Ballesteros-Weinstein numberings from the GPCRdb (gpcrdb.org). As the Ballesteros-Weinstein numbering scheme does not transfer cleanly between different GPCR families, only rhodopsin-like GPCRs were used for distance measurement (11 structures for arrestins, 171 for Gα, and 2 for GRKs; see all distances measured in ‘Distance measurements.xlsx’). Distances between the Cβ carbons on GPCRs and transducers or GRKs <9.0Å were considered residue pairs likely to form an interaction. The determination of the five residues within the intracellular cavity most likely to engage with the two leucines on the transducers and the three aliphatic residues on GRKs (3.54, 5.65, 6.33, 6.36, and 6.37) was based on distance measurements as well as their conserved hydrophobic nature. Mutation of these residues in experimentally solved or AlphaFold3 generated structural models of β2AR, 5-HT_2B_R, and NTR1 was performed manually in PyMOL. The BSA surrounding the residues of the βarr1-FL, Gα-H5, and GRK2-H1 residues when bound to the WT and quintuple mutant GPCRs was determined by *Protein Interfaces, Surfaces and Assemblies* (*PISA*) server (ebi.ac.uk/pdbe/pisa/).

### Bioinformatic analyses

Sequence alignments of the C-terminal of human Gα subunits, the N-terminal of human GRKs, or the finger loop region of human arrestins were performed using *Clustal Omega* (ebi.ac.uk/jdispatcher/msa/clustalo). Logo of the conserved residue plot illustrating the common residues at key position within the intracellular cavity was generated from *WebLogo* server (weblogo.berkley.edu).

## Data analysis

All graphs were generated and analyzed using GraphPad Prism 10 (GraphPad Software). Data are presented as mean ± SEM and N refers to the number of independent experiments (or biological replicates) that have been conducted. Differences were assessed using Student’s *t*-test for two comparisons and one-or two-way ANOVA and Tukey’s post hoc test for multiple comparisons. p<0.05 was considered significant at the 95% confidence level.

## Acknowledgements

We thank Dr. Asuka Inoue for the generous gifts of the knock-out cell lines (βarr1/2 double K.O. and Gα total K.O. cell lines). We are also grateful to Drs. Harsh Bansia and Amedee des Georges for druitful discussions. This work received support from the LEO Foundation (LF18043 to ARBT), NIH grants (R35GM147088 (NIGMS) and R21CA243052 (NCI) to ARBT), and a BBSRC New Investigator Award (BB/X002578/1 to BP).

## Author contributions

H.H. and A.R.B.T. conceived the study; H.H., A.N., and A.R.B.T. designed experiments; H.H., E.F.-E., A.N., and A.R.B.T. conducted experiments and analyzed data; H.H., A.N., M.J., B.P., and A.R.B.T. contributed to the methodology; H.H., and A.R.B.T. drafted the original manuscript with input from all authors; A.R.B.T. supervised the project.

## Materials & Correspondence

Further information and requests for resources and reagents should be directed to and will be fulfilled by the lead contact, Alex R. B. Thomsen. All plasmid constructs and cell lines generated by the authors will be distributed upon request.

---

**Supplementary Figure 1.**
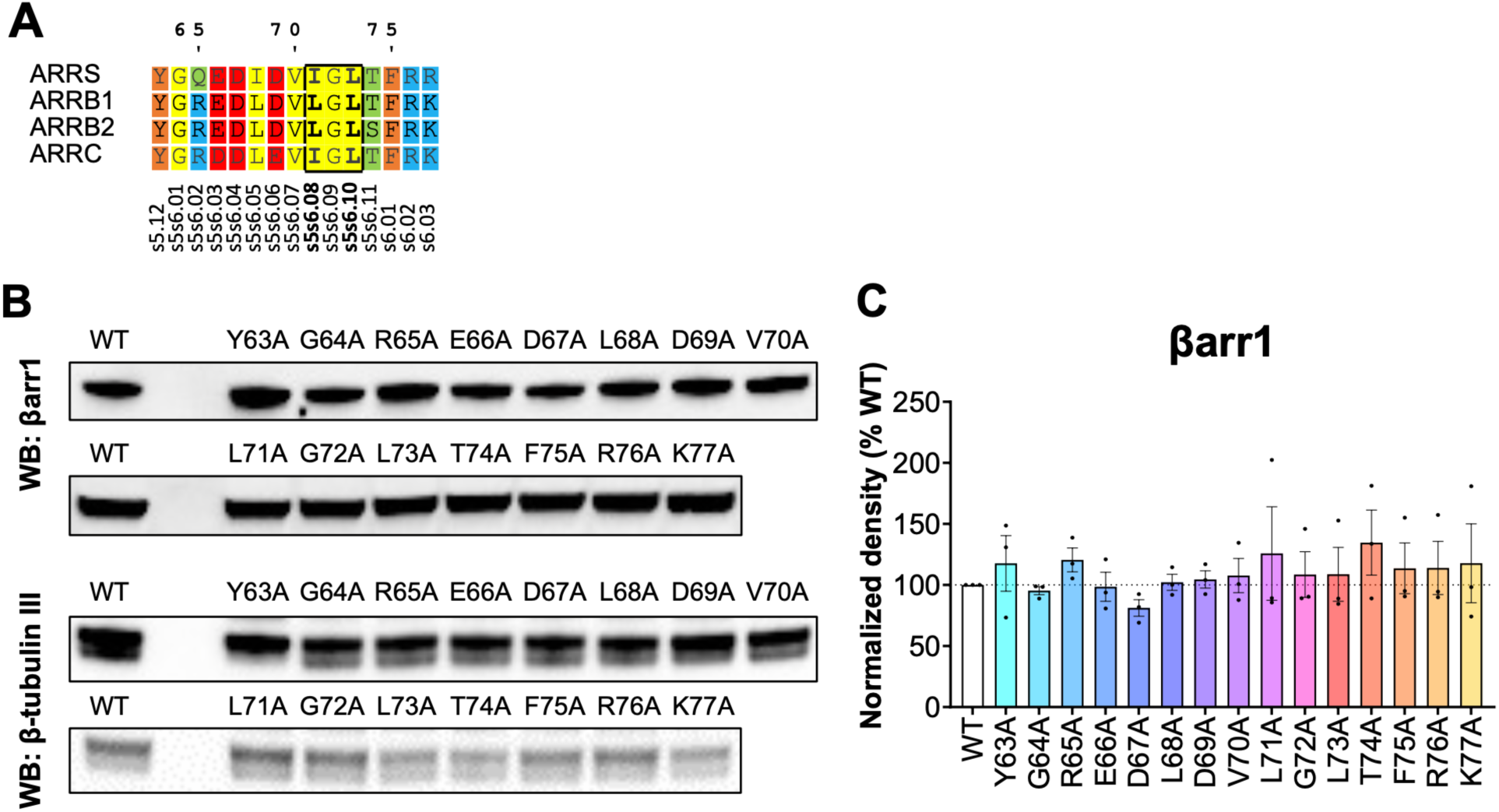
Multiple sequence alignment of arrestins (arrs) and protein expression of βarr1 mutants. (**A**) Multiple sequence alignment of the finger loop residues from arr paralogs in human. βarr1 residue numbering (top) and generic residue numbering (bottom) are shown. Conserved aliphatic residues corresponding to βarr1-L71/L73 are shown in bold. (**B**) Western blots of wild type (WT) and mutants βarr1-SmBiT expression using anti-βarr1 (A1CT) antibody or anti-β-tubulin III antibody. (**C**) Quantification of βarr1-SmBiT protein expression normalized to the β-tubulin III expression. Data represent the means ± SEM of N=3 independent experiments, and one-way ANOVA with Dunnett’s multiple comparisons post hoc test was performed to determine statistical differences between βarr1-SmBiT WT and mutants.

**Supplementary Figure 2.**
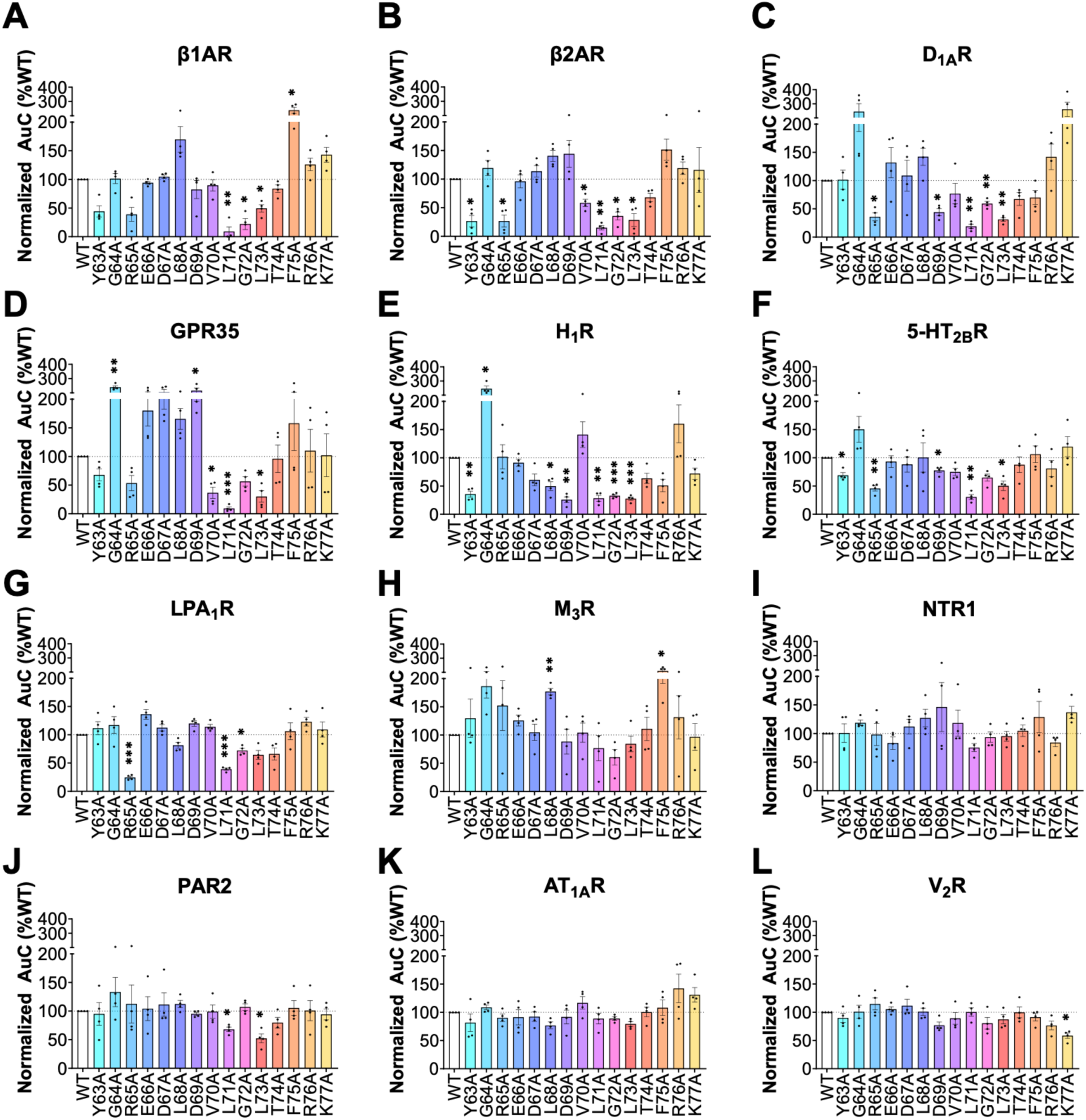
The area under the curve (AuC) quantification associated with results displayed in Fig. 1E is plotted as bar graphs. Data represent the means ± SEM of N=3–4 independent experiments, and one-way ANOVA with Dunnett’s multiple comparisons post hoc test was performed to determine statistical differences between βarr1-SmBiT WT and mutants for each GPCR (\**p* < 0.0332; \*\**p* < 0.0021; \*\*\**p* < 0.0002).

**Supplementary Figure 3.**
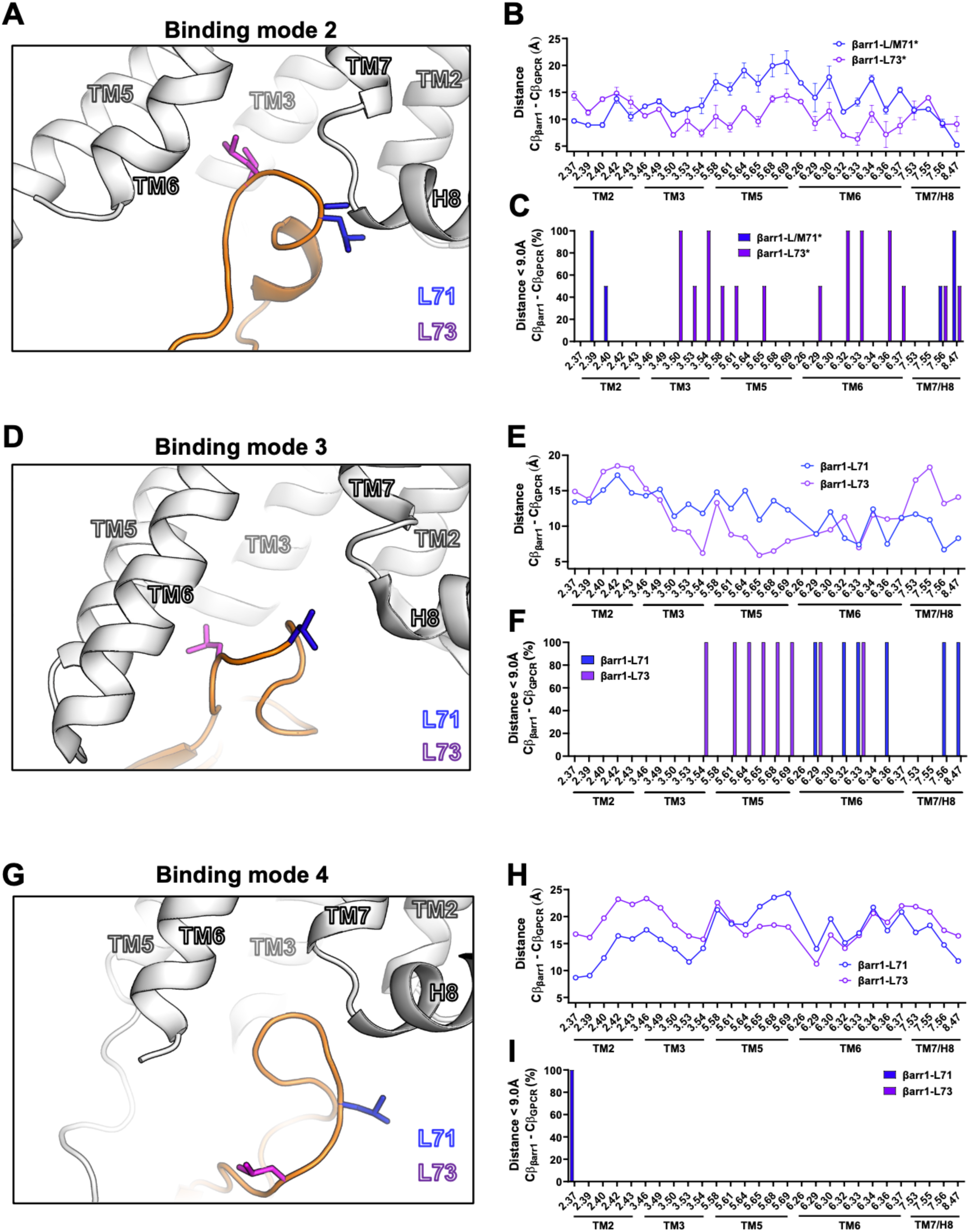
(**A**, **D**, and **G**) Example of GPCR–βarr1 structures (adopted from PDB ID (**A**) 8WRZ, (**D**), 9KUX, and (**G**) 7R9C) showing the relative position of the leucine βarr1-L71/L73 in respect to the receptor in (**A**) *binding mode 2*, (**D**) *binding mode 3*, and (**G**) *binding mode 4*. (**B**, **E**, and **H**) Distance plot quantifying the average distances ± SD between Cβ carbon of βarr1-L/M71 (methionine fr bovine visual rhodopsin) or βarr1-L73 and Cβ carbons of all residues within the GPCR intracellular cavity in (**B**) *binding mode 2*, (**E**) *binding mode 3*, and (**H**) *binding mode 4*. (**C, F**, and **I**) Bar graph showing the percentage of structures where the distance between Cβ carbon of βarr1-L/M71 or βarr1-L73 and each individual Cβ carbons of all residues within the GPCR intracellular cavity is below 9.0Å in (**C**) *binding mode 2*, (**F**) *binding mode 3*, and (**I**) *binding mode 4*.

**Supplementary Figure 4.**
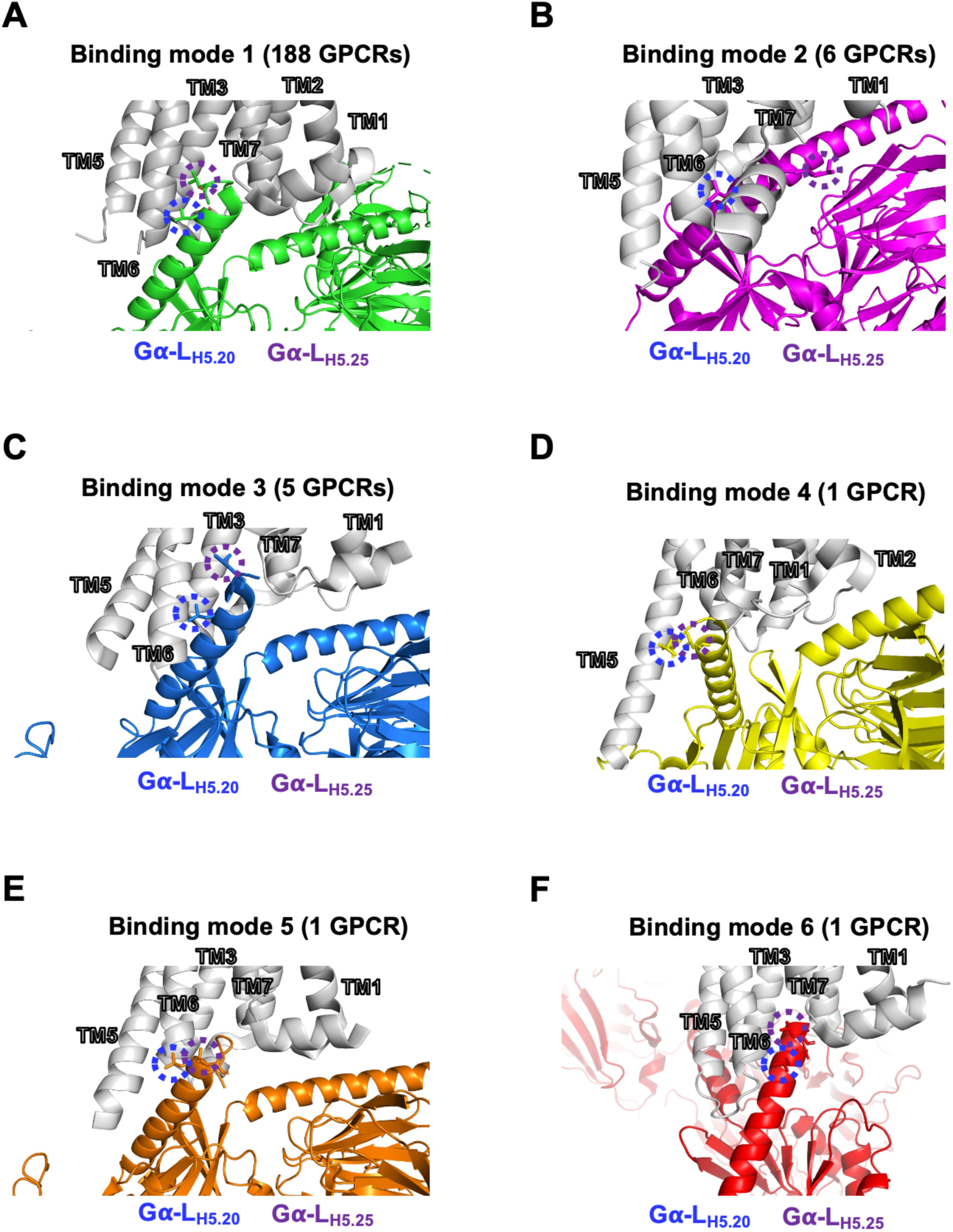
Example of GPCR–G protein structure showing the relative position of the leucine residues at H5.20 and H5.25 in respect to the receptor in (**A**) *binding mode 1* (based on 7SRR), (**B**) *binding mode 2* (based on 7CX2), (**C**) *binding mode 3* (based on 7XOV), (**D**) *binding mode 4* (based on 8HMP), (**E**) *binding mode 5* (based on 8K8J), and (**F**) *binding mode 6* (based on 7RKN).

**Supplementary Figure 5.**
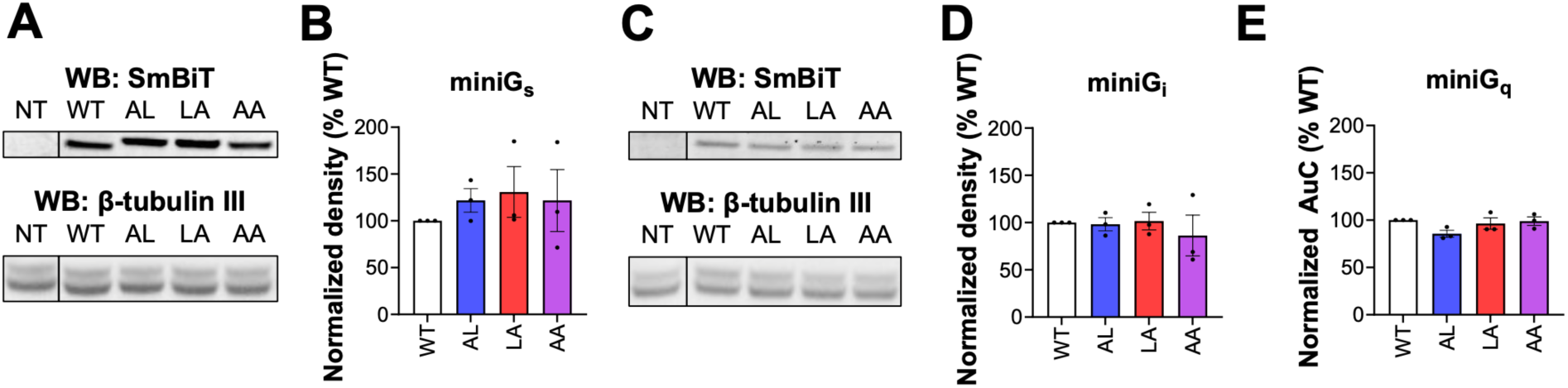
Quantification of miniG-SmBiT expression by western blots. (**A**) Western blots of wild type (WT) and mutants (AL, LA, and AA) SmBiT-miniG_s_ using anti-SmBiT or anti-β-tubulin III antibodies. (**B**) Quantification of miniG_s_-SmBiT expression normalized to the β-tubulin III expression. (**C**) Western blots of wild type (WT) and mutants (AL, LA, and AA) SmBiT-miniG_i_ using anti-SmBiT or anti-β-tubulin III antibodies. (**D**) Quantification of miniG_i_-SmBiT expression normalized to the β-tubulin III expression. (**E**) Quantification of wild type (WT) and mutants (AL, LA, and AA) SmBiT-miniG_q_ relative expression using baseline luminescence generated by random collision with cytosolic-expressed LgBiT in the NanoBiT assay (quantified in this way due to very low/non-detectable expression by western blotting).

**Supplementary Figure 6.**
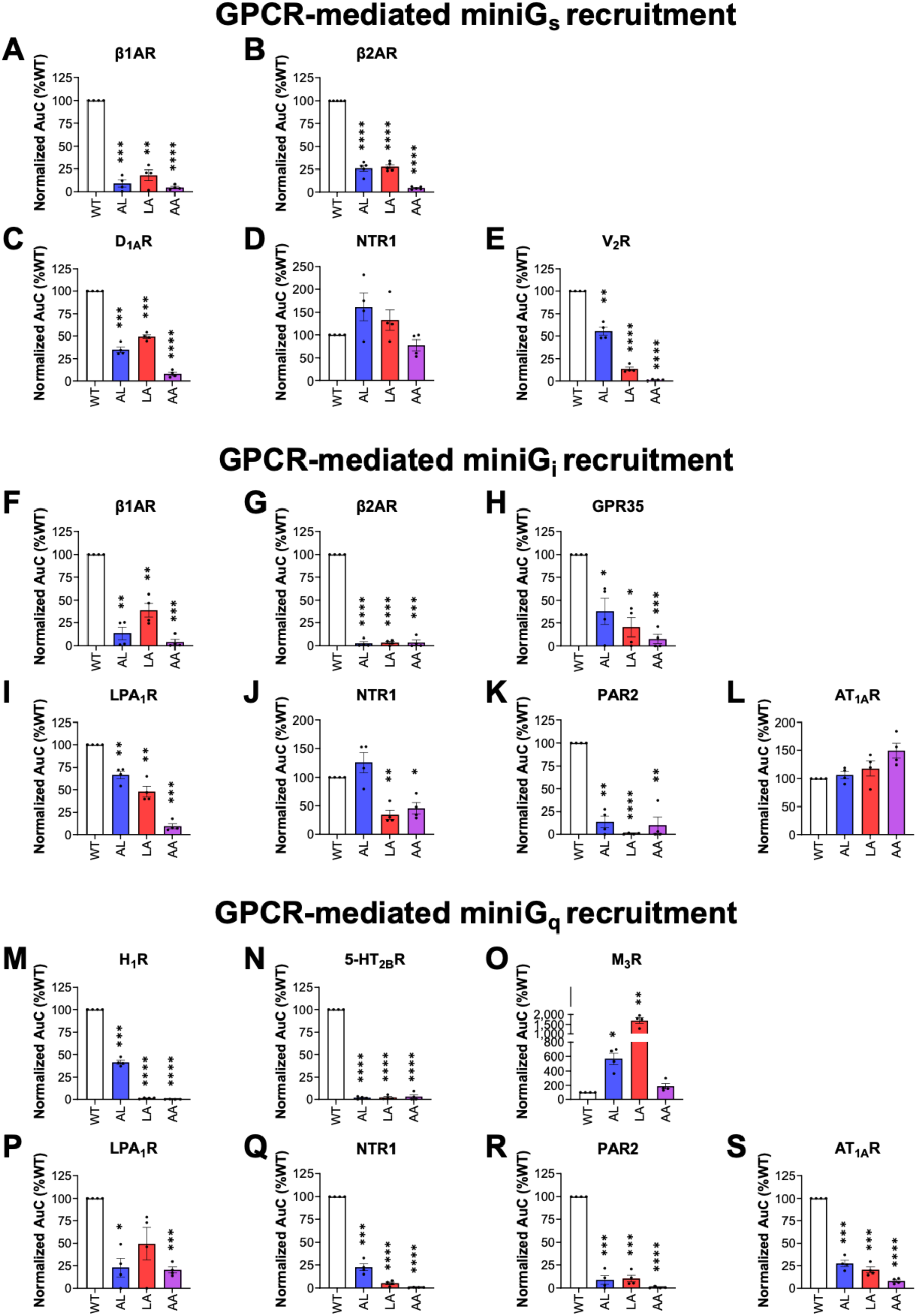
Importance of the Gα subunit C-terminal L_H5.20_ and L_H5.25_ residues for agonist-stimulated GPCR-G protein interaction. (**A**-**S**) The area under the curve (AuC) quantification associated with results displayed in (**A**-**E**) Fig. 4H, (**F**-**L**) Fig. 4I, and (**M**-**S**) Fig. 4J is plotted as bar graphs. (**A**-**S**) Data represent the means ± SEM of N=4 independent experiments, and one-way ANOVA with Dunnett’s multiple comparisons post hoc test was performed to determine statistical differences between agonist-stimulated cells transfected with WT miniG proteins and mutants (\**p*<0.0322 \*\**p* < 0.0021; \*\*\**p* < 0.0002; \*\*\*\**p*<0.0001).

**Supplementary Figure 7.**
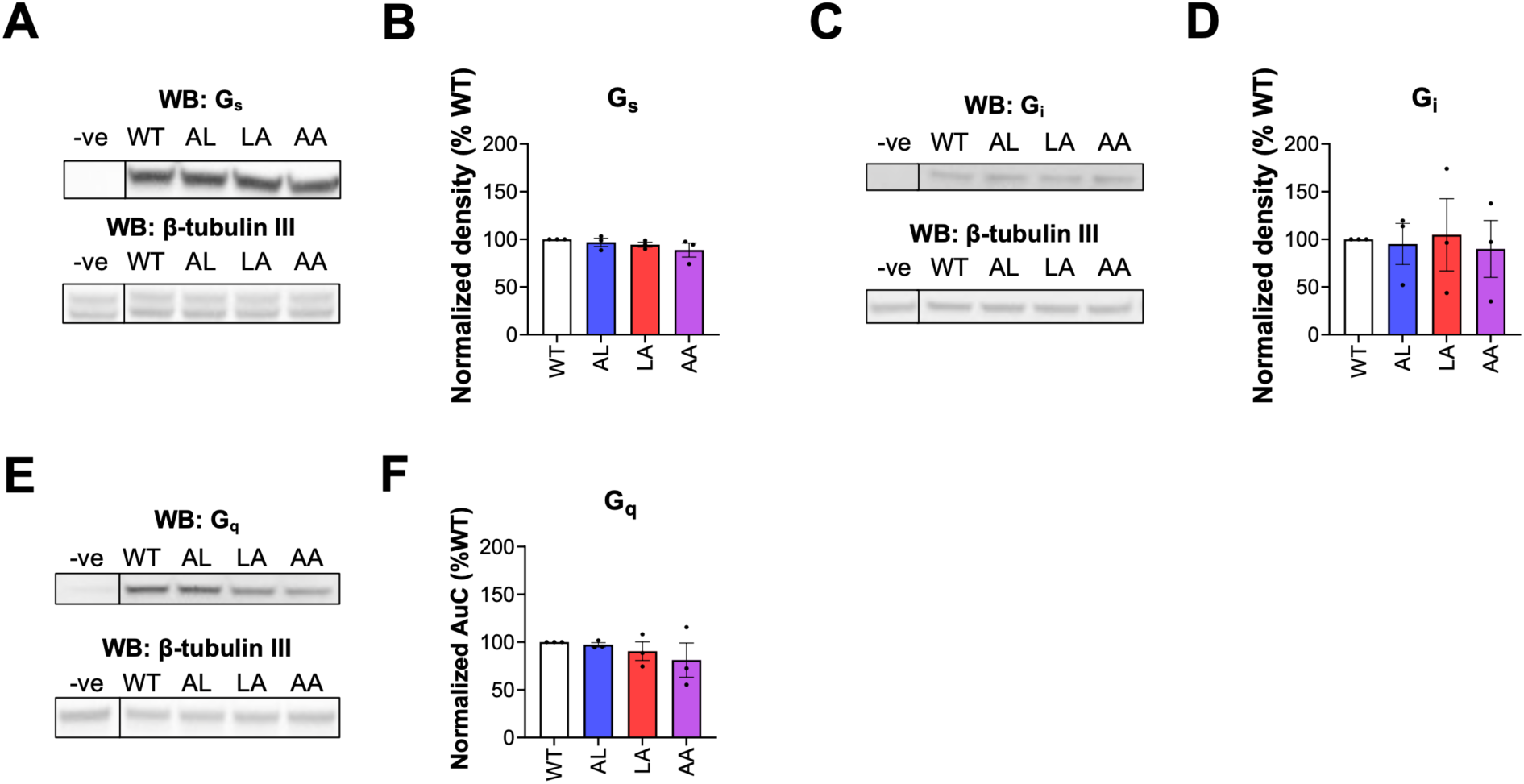
Protein expression of Gα subunit in Gα total knock-out HEK293 cells by western blotting. (**A**) Western blots of Gα_s_ wild type (WT) and mutants (AL, LA, and AA) using anti-Gα_s_ or anti-β-tubulin III antibodies. (**B**) Quantification of Gα_s_ protein expression normalized to the β-tubulin III expression. (**C**) Western blots of Gα_i1_ wild type (WT) and mutants (AL, LA, and AA) using anti-Gα_i1_ or anti-β-tubulin III antibodies. (**D**) Quantification of Gα_i1_ protein expression normalized to the β-tubulin III expression. (**E**) Western blots of Gα_q_ wild type (WT) and mutants (AL, LA, and AA) using anti-Gα_q_ or anti-β-tubulin III antibodies. (**F**) Quantification of Gα_q_ protein expression normalized to the β-tubulin III expression.

**Supplementary Figure 8.**
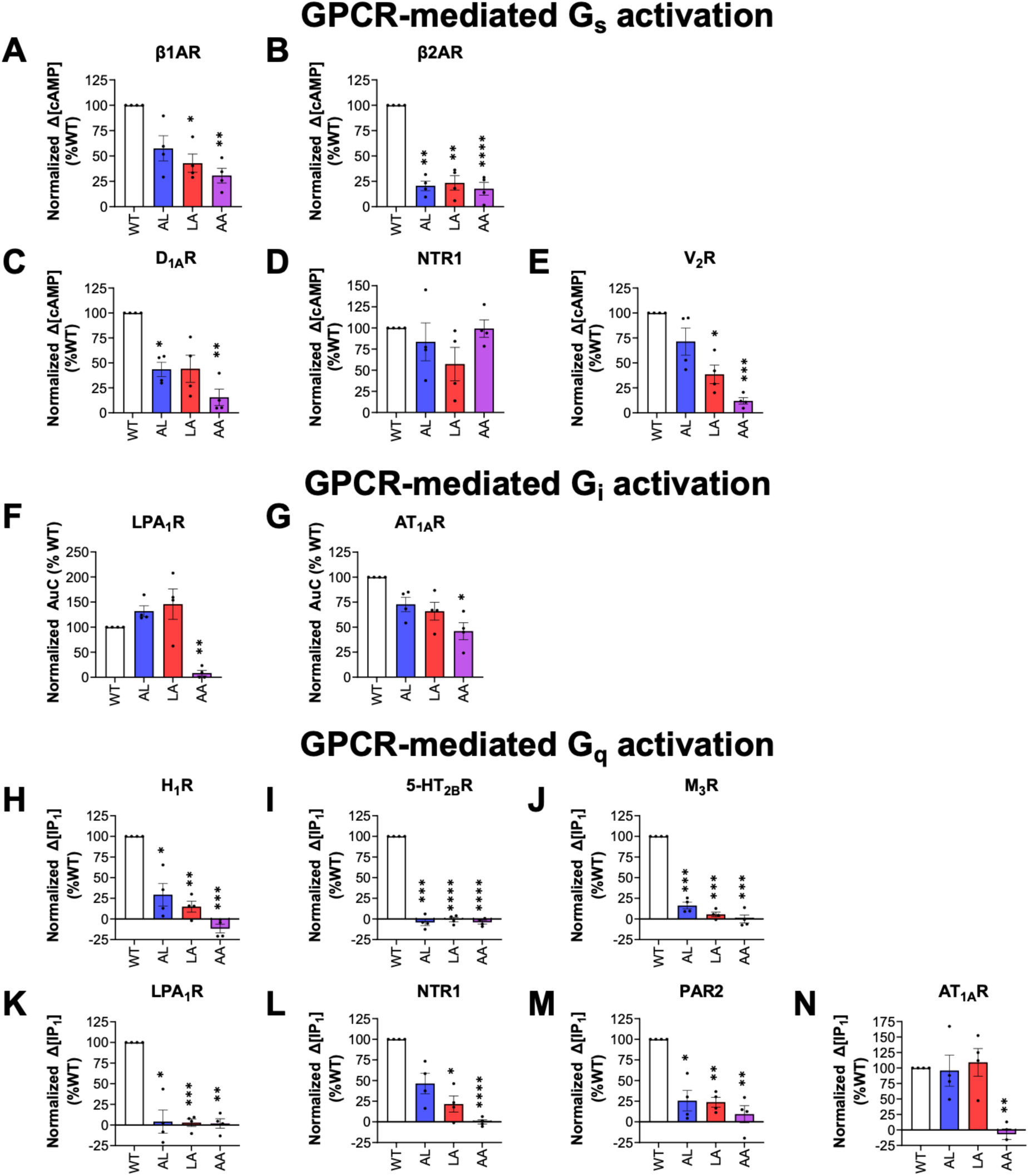
Importance of the Gα subunit C-terminal L_H5.20_ and L_H5.25_ residues for agonist-stimulated G protein activation. (**A**-**E**) Agonist-stimulated cAMP production in HEK293 total Gα K.O. cells transfected with G_s_-coupled GPCRs and WT or mutants Gα_s_. (**F**-**G**) Agonist-stimulated G_i_ activation in HEK293 total Gα K.O. cells transfected with G_i/o_-coupled GPCRs and WT or mutants Gα_i1_. Gα_i1_ activation was measured by NanoBiT assay between GPCR-LgBiT and Rap1GAP-SmBiT. (**H**-**N**) Agonist-stimulated IP_1_ production in HEK293 total Gα K.O. cells transfected with G_q/11_-coupled GPCRs and WT or mutants Gα_q_. (**A**-**N**) Data represent the means ± SEM of N=4 independent experiments, and one-way ANOVA with Dunnett’s multiple comparisons post hoc test was performed to determine statistical differences between agonist-stimulated cells transfected with WT Gα subunit and mutant Gα subunits (\**p*<0.0322 \*\**p* < 0.0021; \*\*\**p* < 0.0002; \*\*\*\**p*<0.0001).

**Supplementary Figure 9.**
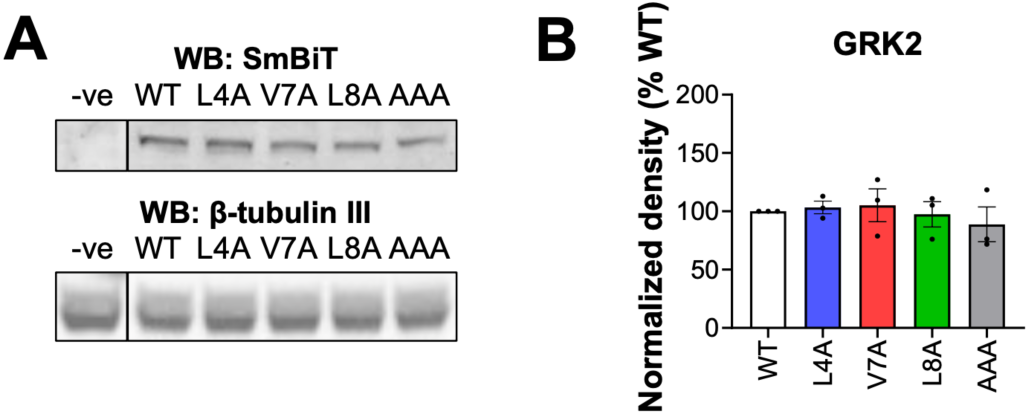
Quantification of GRK2 expression in HEK293 cells by western blots. (**A**) Western blots of wild type (WT) and mutants (L4A, V7A, L8A, and triple AAA mutant) GRK2 using anti-GRK2 or anti-β-tubulin III antibodies. (**B**) Quantification of GRK2 expression normalized to the β-tubulin III expression.

**Supplementary Figure 10.**
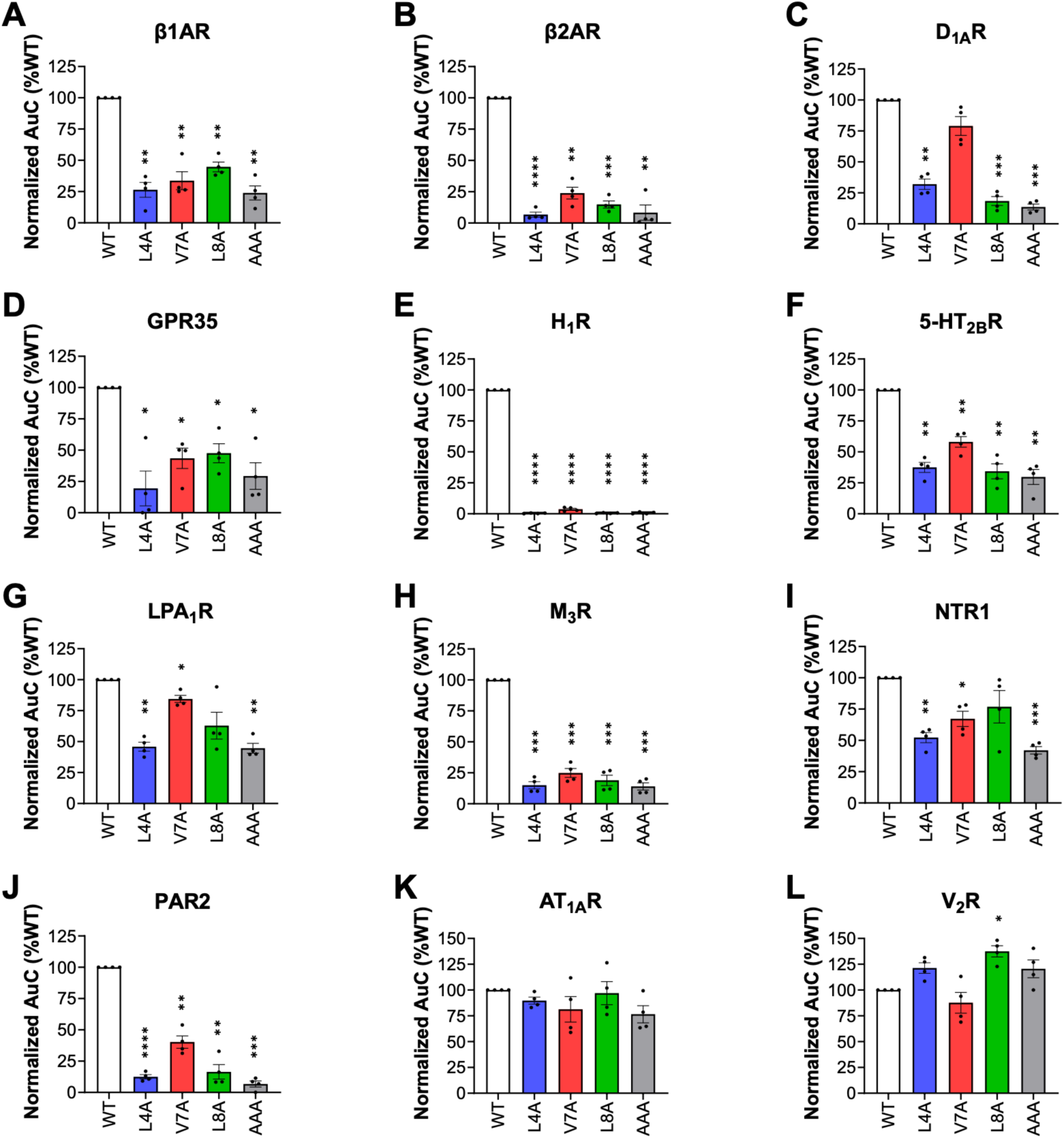
The area under the curve (AuC) quantification associated with results displayed in Fig. 5J is plotted as bar graphs. Data represent the means ± SEM of N=4 independent experiments, and one-way ANOVA with Dunnett’s multiple comparisons post hoc test was performed to determine statistical differences between agonist-stimulated cells transfected with WT GRK2 and mutants (\**p*<0.0322 \*\**p* < 0.0021; \*\*\**p* < 0.0002; \*\*\*\**p*<0.0001).

**Supplementary Figure 11.**
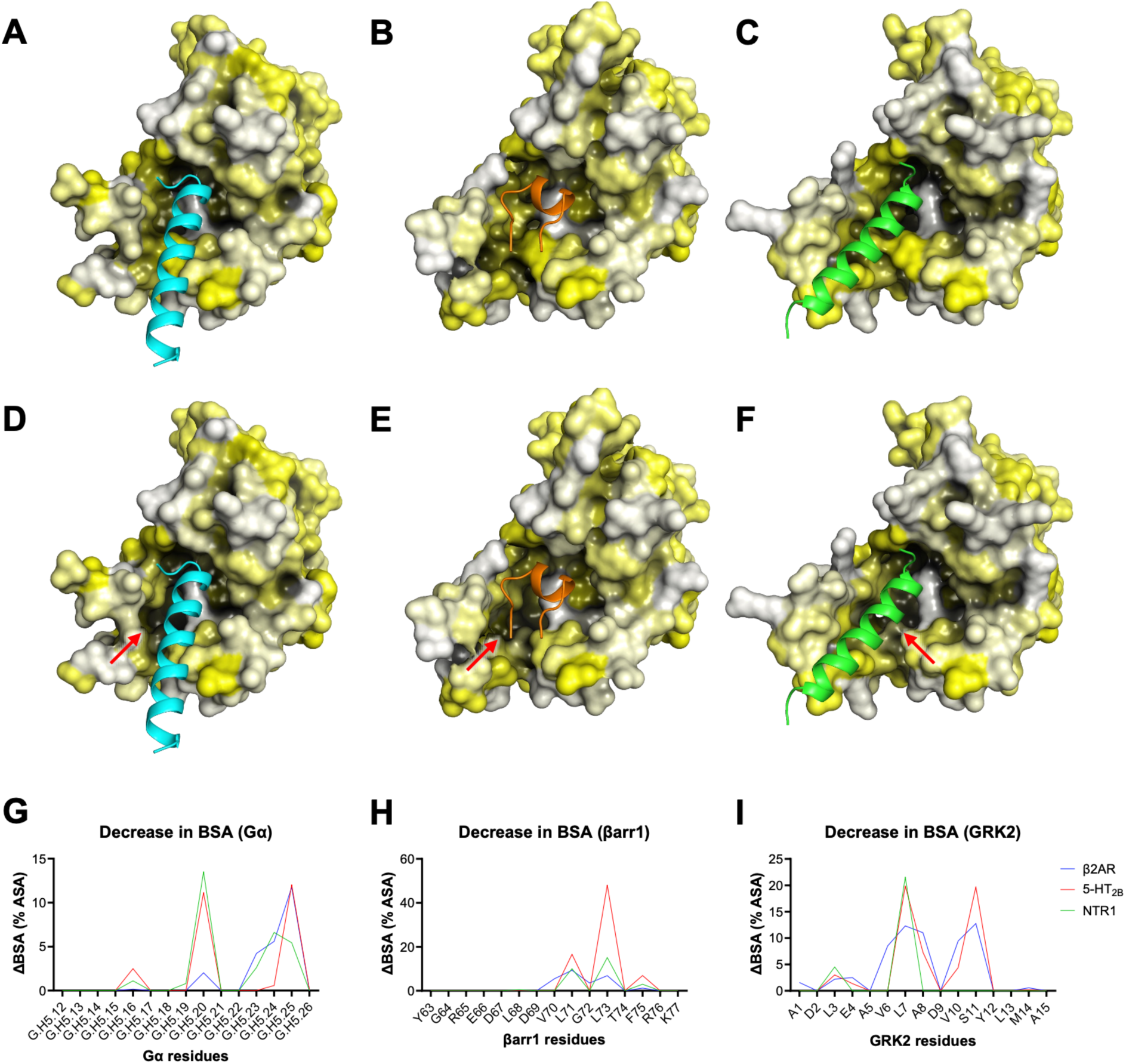
Decrease in hydrophobic patch surface area upon computational introduction of the quintuple mutations. (**A**–**C**) Surface representations of 5-HT_2B_R in complex with (**A**) miniG_q_ (PDB ID: 7SRR), (**B**) βarr1 (PDB ID: 7SRS), or (**C**) GRK2 (modeled with AlphaFold3). (**D**–**F**) Surface representations of 5-HT_2B_R harboring the mutations at positions 3.54, 5.65, 6.33, 6.36, and 6.37 (AAGAA), in complex with (**D**) miniG_q_, (**E**) βarr1, or (**F**) GRK2, show the partial loss of the hydrophobic patch at the interface with the transducers. (**G**–**I**) Quantification of decrease in buried surface area (BSA) around each residue on (**G**) Gα, (**H**) βarr1, or (**I**) GRK2, at their interfaces with β2AR, 5-HT_2B_R, or NTR1. Introducing the quintuple mutations to the GPCRs exposes the key aliphatic residues on the transducers (G.H5.20 and G.H5.25 on Gα, L71 and L73 on βarr1, and L3, V6, and L7 on GRK2) as indicated by the decrease in BSA.

**Supplementary Figure 12.**
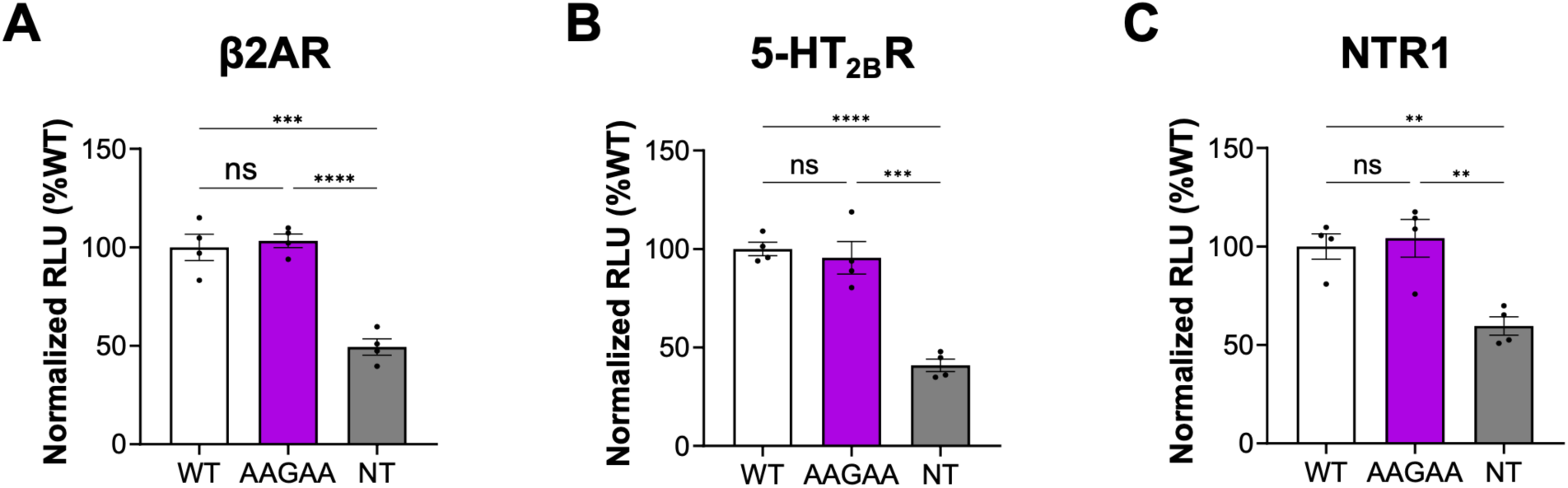
Quantification of GPCR surface expression in HEK293 cells transfected with either wild-type, AAGAA mutant, or mock of (**A**) β2AR, (**B**) 5-HT_2B_R, and (**C**) NTR1 by ELISA using an anti-FLAG antibody. Data represents the means ± SEM of N=4 independent experiments, and one-way ANOVA with Dunnett’s multiple comparisons post hoc test was performed to determine statistical differences between mock transfected (NT) cells and each GPCR constructs (\*\**p* < 0.0021; \*\*\**p* < 0.0002; \*\*\*\**p*<0.0001).

